# SOD1 Catalyses Thiol Oxidation to Thiosulfinates

**DOI:** 10.64898/2026.06.03.729904

**Authors:** Christopher H. Switzer

## Abstract

Cu/Zn superoxide dismutase (SOD1) is canonically regarded as a superoxide scavenging antioxidant yet is paradoxically associated with multiple diseases. Here, we show that SOD1 catalyzes thiol oxidation to thiosulfinates (RS(O)SR), revealing a previously unrecognized copper mediated oxidant forming reaction in biology. Steady state kinetics demonstrate robust, O_2_ dependent thiol consumption, and ATR FTIR spectra of the SOD1–cysteine reaction display S–O bands (≈1042/1169 cm^−1^) identical to cysteine thiosulfinate, including matching pseudo–first order decay in excess cysteine. SOD1 generated thiosulfinates are potent electrophiles and oxidants, supported by dimedone trapping and O-atom transfer to TCEP and horseradish peroxidase, with alkaline lability consistent with thiosulfinate hydrolysis rather than H_2_O_2_. Exogenous thiosulfinates phenocopy SOD1 mediated thiol oxidation, including GSH depletion, protein sulfenylation and cytotoxicity. SOD1 inhibition strongly suppresses cysteine and homocysteine induced toxicity, demonstrating that thiol driven oxidative stress requires SOD1 activity. SOD1 overexpression in human cells triggers oxidative stress and reduces proliferation, an effect that is absent in a copper-deficient mutant. N-acetylcysteine treatment further amplified this SOD1-dependent oxidative stress. At lower levels, nanomolar thiosulfinates elicit a hormetic proliferative response and rescue SOD1 deficient growth, identifying a pro-growth signaling function mediated by basal thiosulfinate formation. Kinetic modelling indicates that thiol–thiosulfinate turnover can match basal superoxide dismutation, indicating that thiosulfinate synthesis is a major catalytic output of SOD1. These findings identify SOD1 as a thiol oxidizing enzyme that generates thiosulfinates, establishing a core sulfur-based oxidation pathway and revealing that two classical “antioxidants”, SOD1 and thiols, together generate potent oxidants that link thiol metabolism to both cytotoxic and growth promoting outcomes.

**Significance Statement:** Although SOD1 is classically defined as an antioxidant, its association with diverse oxidative-stress–driven diseases suggests additional chemistry at work. Here we identify thiosulfinate synthesis as a major catalytic output of SOD1, revealing that the enzyme is not merely a superoxide detoxifier but a thiol-oxidizing catalyst. This activity provides a unifying mechanism for two long-standing biological paradoxes: the unexplained toxicity of elevated thiols and the pro-growth, pro-survival signaling linked to basal SOD1 activity. By establishing thiosulfinates as a central oxidative currency in cells, this work reframes SOD1 as a bifunctional oxidase that shapes both stress responses and proliferative programs.

## Introduction

Superoxide dismutase 1 (SOD1) is a ubiquitous and abundant Cu/Zn enzyme that catalyzes superoxide (O_2_^−^) detoxification and plays a central role in cellular ROS metabolism (1). Its high expression levels and antioxidant activity have long supported the view of SOD1 as a broadly protective redox enzyme (2). Yet elevated or dysregulated SOD1 activity is frequently maladaptive in disease, suggesting that the enzyme’s chemistry may extend beyond classical ROS detoxification (2-7). The solvent-exposed copper center provides access to additional redox pathways, and SOD1 oxidizes hydrogen sulfide (H_2_S) (8, 9) and several low-molecular-weight thiols (10-12), indicating that the enzyme participates in broader sulfur-based chemistry than previously appreciated. Notably, early reports that SOD1 and thiols generate H_2_O_2_ lacked a defined chemical mechanism (10, 11), leaving the identity of the oxidant produced during SOD1-mediated thiol turnover unresolved.

Cellular thiols such as cysteine and homocysteine are indispensable for redox balance and core biosynthetic pathways. Cysteine is widely regarded as an antioxidant because it fuels glutathione synthesis and can directly quench electrophiles and oxidants. Yet, despite this protective reputation, elevated thiol levels are cytotoxic: cysteine induces rapid loss of viability in mammalian cells and animal models (13-16), millimolar cysteine is lethal to cultured human cells (14), and chronic N-acetylcysteine administration promotes lung adenocarcinomas in mice (17). Homocysteine, strongly associated with cardiometabolic and neurological disease, similarly triggers oxidative stress and cell death when elevated (18-20). How molecules considered “antioxidants” can produce immediate and extensive oxidative damage remains an unresolved paradox in redox biology (14, 16, 21, 22). This disconnect points to a missing chemical step linking thiol abundance to oxidative outcomes.

Here, thiosulfinates (RS(O)SR) are identified as the direct products of SOD1-catalysed thiol oxidation, establishing that SOD1 functions as a cellular thiol oxidase capable of generating sulfur-based oxidants/electrophiles. Spectroscopic, kinetic, and reactivity data demonstrate that these thiosulfinates arise during SOD1 turnover with small thiols and rapidly engage in oxygen-atom transfer and sulfenylation reactions characteristic of electrophilic sulfur oxidants. Because thiosulfinates react readily with cellular thiols (23-26), their formation provides a unifying chemical framework linking thiol abundance to divergent biological outcomes: at high thiol flux, SOD1-derived thiosulfinates drive glutathione loss, protein sulfenylation, and cytotoxicity, whereas at basal flux, the same chemistry produces a hormetic, pro-growth signal. Most strikingly, these findings reveal that SOD1 converts abundant cellular thiols into a sulfur-based oxidant, establishing an unrecognized oxidative output of SOD1 that operates alongside its classical superoxide metabolizing function.

## Results

### SOD1 catalyzes thiol oxidation to form an oxidant

Thiol oxidation was quantified by incubating SOD1 with increasing L-cysteine and monitoring loss of free RSH over 30 min (Fig. 1A). Global fitting of steady-state initial rates to the Michaelis–Menten equation yielded V_max_ = 356.3 µM·min^−1^ and K_M_ = 3.57 mM for cysteine consumption (Fig. 1B and SI Appendix, Table S1). With E_t_ = 10 µM, the corresponding turnover number is k_cat_ = 35.63 min^−1^ (0.594 s^−1^), and the specificity constant is k_cat_/K_M_ = 3.32 × 10^2^ M^−1^·s^−1^. Because K_M_ greatly exceeds intracellular free cysteine, SOD1 functions in the [S] ≪ K_M_ condition in cells, where v ≈ (k_cat_/K_M_)[E_t_][S]; accordingly, initial rates scale linearly with micromolar cysteine (Fig. 1C). Additionally, D/L-homocysteine exhibited kinetics similar to cysteine, while dihydro-lipoic acid was oxidized at reduced rates (Fig. 1B). Glutathione (GSH) showed no measurable SOD1-dependent oxidation. Concomitant oxidant formation during turnover was verified using coumarin boronic acid (CBA) fluorescence, which increased with time and cysteine concentration (Fig. 1D). CBA oxidation rates mirrored thiol-oxidation kinetics for cysteine and dihydro-lipoic acid (Fig. 1E). Furthermore, CBA oxidation rates correlated linearly with SOD1 concentration (Fig. 1F). Oxidant production requires O_2_, as increasing concentrations of SOD1 did not significantly increase CBA fluorescence in hypoxic conditions. The observed O_2_-dependent thiol consumption and concomitant oxidant production confirm that SOD1 carries intrinsic thiol-oxidizing activity, aligning with earlier reports (10).

**Fig. 1.**
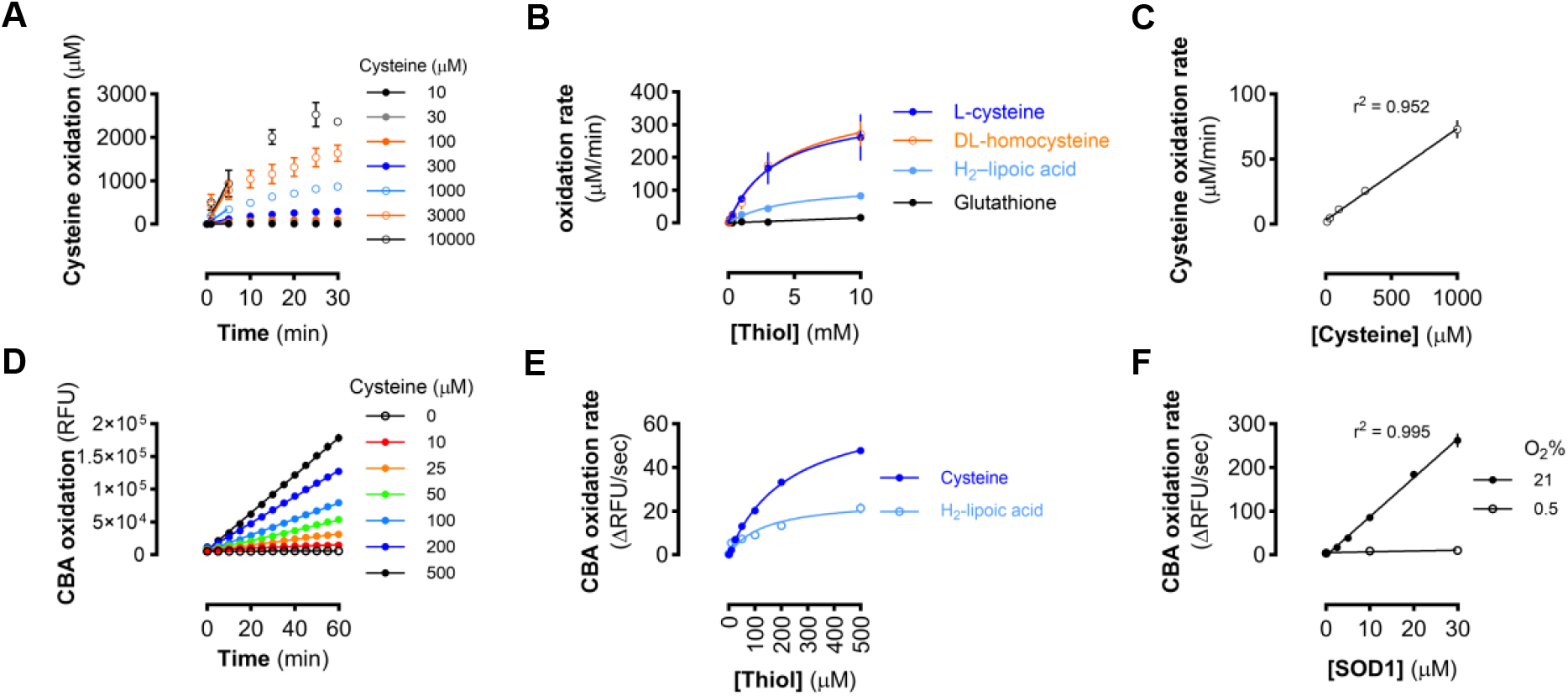
SOD1 catalyzes thiol oxidation and generates an oxidant. (A) Cysteine consumption over time for increasing cysteine concentrations in the presence of SOD1. Data are mean ± SEM (n = 3). Initial rates were determined by linear regression analysis. (B) Oxidation rates of thiols in the presence of SOD1. Data are mean ± SEM (n = 3-6) and fit to Michaelis-Menten equation (Supporting Information, Table S1). (C) Linear increase in cysteine-oxidation rate at sub-millimolar cysteine. Points show mean ± SEM (n = 4-6) and were fitted by linear regression (solid line; r^2^ = 0.952). (D) CBA oxidation over time for increasing cysteine concentrations in the presence of SOD1. Data are mean ± SEM (n = 4). (E) Oxidation rates of CBA in the presence of SOD1 and cysteine or dihydro-lipoic acid. Data are mean ± SEM (n = 3-6) and fit to pseudo-first order kinetics (r^2^_(Cys)_ = 0.993; r^2^_(H2-LA)_ = 0.805). (F) CBA oxidation rates in response to cysteine (300 µM) and varying SOD1 concentration in 21% or 0.5% O_2_ atmosphere. Data are mean ± SEM (n = 3) and analysed by linear regression (r^2^_(21%)_ = 0.995; r^2^_(0.5%)_ = 0.791).

SOD1-catalysed sulfide oxidation is proposed to proceed through one-electron transfer from HS^−^ to Cu(II), generating a sulfanyl radical (HS•) (8). By analogy, SOD1-mediated thiol oxidation is expected to form [Cu–SR]+ thiyl-like intermediates. Oxidant production from this initial intermediate could proceed via two pathways: (1) disulfide radical anion formation (via attack by a second thiolate) followed by O_2_ reduction, or (2) O_2_ insertion into the [Cu–SR]^+^ species to give a Cu(II)– sulfinate (SI Appendix, Fig. S1A). One pathway proceeds through a high-energy disulfide radical anion, whereas the other reaction pathway yields the thiosulfinate after nucleophilic attack by another thiolate. To determine if a disulfide radical anion is integral to oxidant formation, ferricyanide was utilized as an alternative electron sink in the SOD1–cysteine reaction (27). CBA oxidation rates were not significantly altered in the presence of ferricyanide (SI Appendix, Fig. S1B), indicating that a high energy reductant is not formed and suggesting that the O_2_ insertion mechanism results in thiosulfinate formation by SOD1.

Thiosulfinates are electrophilic species characterized by a relatively weak S–S bond that undergoes rapid alkaline hydrolysis to the corresponding sulfenic acid (RSOH).(26) To determine whether cysteine thiosulfinate (CTS) is the enzymatic product formed when cysteine is used as substrate, the pH dependence of cysteine consumption and oxidant generation was examined. Cysteine consumption increased with rising pH (Fig. 2A). In contrast, CBA oxidation displayed a distinct bell-shaped pH profile, with maximal reactivity at pH 7.5 and marked reduction at both lower and higher pH values (Fig. 2A). The loss of detectable oxidant formation under alkaline conditions aligns with the expected base catalyzed hydrolysis of RS(O)SR to RSOH. In contrast, the CBA oxidation rate by H_2_O_2_ was slightly elevated at pH 10 compared to 7.5 (Fig. 2B). An alternative pathway in which *in situ* H_2_O_2_ oxidizes cystine to CTS is unlikely under our conditions as disulfide oxidation by H_2_O_2_ to thiosulfinate requires acidic conditions and proceeds poorly at neutral pH (28), further supporting direct thiosulfinate formation.

**Fig. 2.**
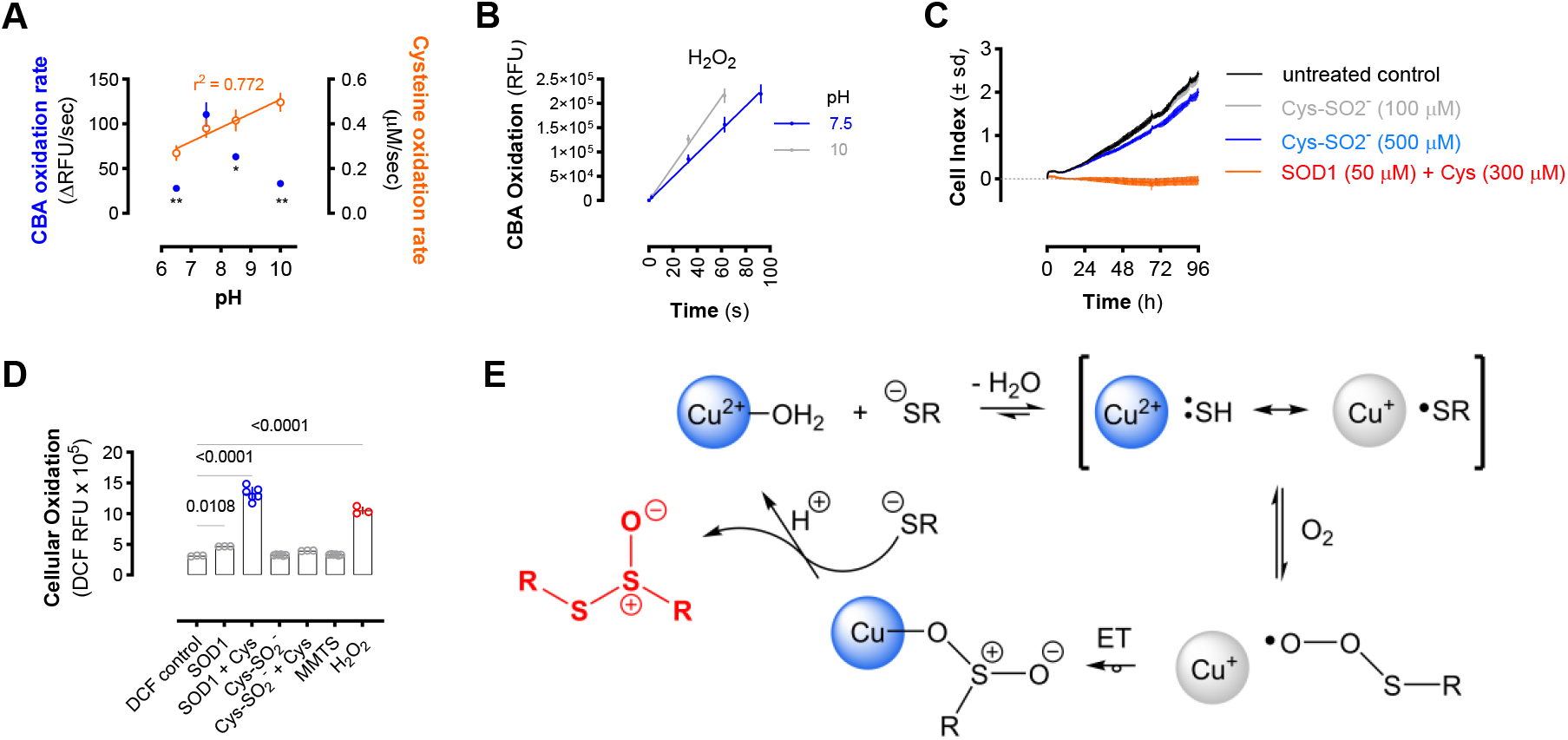
SOD1™thiol reaction forms a thiosulfinate-like oxidant. (A) pH profile of CBA oxidation and cysteine oxidation rates of SOD1™cysteine reaction. Data are mean ± SEM (n = 3-5). CBA oxidation rate (blue) change significance relative to pH 7.5 was calculated by one-way ANOVA with Dunnett’s test and cysteine oxidation rates fitted to linear regression (r^2^ = 0.772). (B) pH profile of H_2_O_2_ mediated CBA oxidation kinetics. H_2_O_2_ oxidised CBA with an elevated rate at pH 10 (grey) compared to pH 7.5 (blue). (C) Cell index (proliferation) of HEK293 cells with cysteine sulfinate (Cys-SO_2_^-^) or SOD1 (50 µM) + cysteine (300 µM) supplemented media and compared to untreated cells. Data are mean cell index values ± sd (n = 4). (D) Cellular oxidation control conditions. HEK293 oxidative stress measured by DCF fluorescence in response to media supplementation with SOD1(50 µM), (50 µM) + cysteine (300 µM), Cys-SO_2_^-^ (300 µM), Cys-SO_2_^-^ + Cysteine (300 µM each), MMTS (300 µM), or H_2_O_2_ (100 µM). Bars are mean DCF RFU values ± sd (n = x) and significance calculated by one-way ANOVA with Dunnett’s test. (E) Proposed mechanism of SOD1 thiosulfinate synthase catalytic cycle. Coordination of thiolate (RS^-^) to Cu(II)-SOD1 forms a [Cu **·**SR]^+^ intermediate. O_2_ insertion forms a sulfanylperoxo that is reduced by Cu(I) and rearranged to a Cu(II) sulfinate. Thiol attack forms the thiosulfinate and starting Cu(II) complex.

Cysteine sulfinic acid (Cys-SO_2_^-^) is a possible product of the SOD1–thiol oxidation pathway; however, exogenous Cys-SO_2_^-^ did not suppress HEK293 proliferation, whereas media containing SOD1 (50 µM) and cysteine (300 µM) completely abolished cell growth (Fig. 2C) (11). Similarly, neither Cys-SO_2_^-^ alone nor in combination with cysteine induced oxidative stress, as assessed by DCF fluorescence (Fig. 2D). Moreover, disulfide stress produced by MMTS also failed to elicit DCF fluorescence (Fig. 2D). Together, these data indicate that neither cysteine sulfinic acid nor disulfide formation accounts for the oxidative signaling, supporting the conclusion that SOD1-dependent thiosulfinate formation is the key oxidant product driving the observed cellular responses (Fig. 2E).

### Spectroscopic and Kinetic Identification of Thiosulfinate Formation

The products of SOD1 cysteine oxidation were analyzed by Fourier transform infrared (FT-IR) spectroscopy. Authentic CTS showed a single, intense ν(S–O) band at 1105 cm^−1^, corresponding to the ν(S–O) stretch of the thiosulfinate functional group (RS(O)–SR) (SI Appendix, Fig. S2A) and a characteristic UV absorbance maximum at 247 nm (SI Appendix, Fig. S2B) (26). In contrast, SOD1 incubated with cysteine produced a broader, evolving S–O envelope centered at ∼1042 cm^-1^ and 1169 cm^-1^ (Fig. 3A), rather than the single CTS peak. Because CTS is expected to react with excess cysteine, FT-IR spectra of CTS + cysteine was acquired under identical conditions, which generated the same two-band pattern centered at ∼1042 cm^−1^ and 1169 cm^−1^ (Fig. 3B). Furthermore, authentic CTS decayed with pseudo–first-order kinetics in the presence of cysteine (k = 1.41×10^-3^ s^-1^; SI Appendix, Fig. S2C), while the 1042 cm^−1^ peak during the SOD1 + cysteine reaction yielded a similar rate constant (k = 1.88×10^-3^ s^-1^; SI Appendix, Fig. S2D). The agreement in spectral features and kinetic behavior indicates that both reactions are governed by the same CTS-driven step, demonstrating that thiosulfinates are direct, transient products of SOD1-mediated thiol oxidation.

**Fig. 3.**
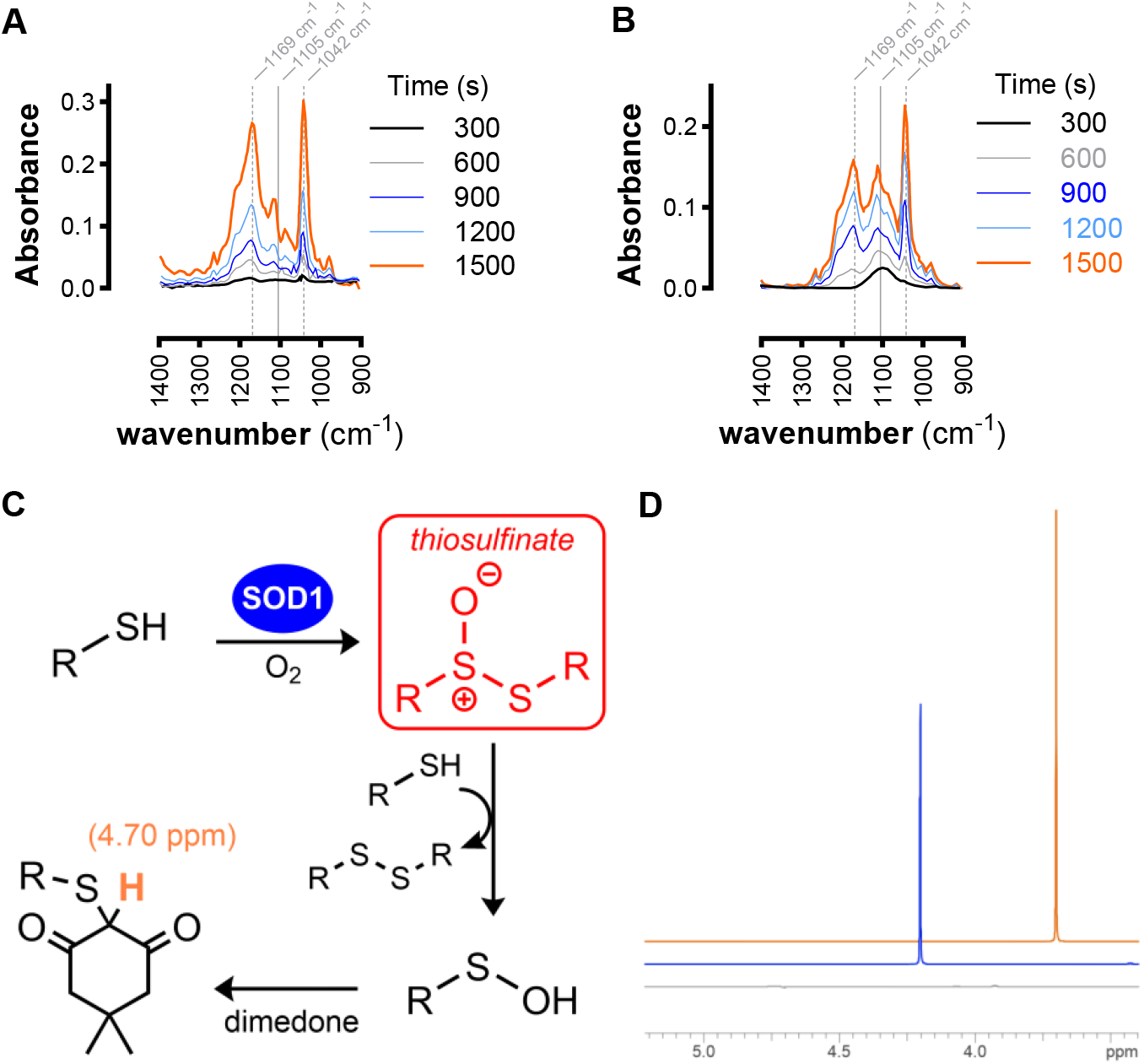
SOD1 generates thiosulfinate during cysteine turnover. (A) Time-resolved FT-IR spectra of SOD1 and (B) CTS reacting with cysteine at pH 7.5. (B) Schematic showing thiol oxidation to thiosulfinate by SOD1 and subsequent reaction with excess thiol to form sulfenic acids, which are trapped by dimedone to form dimedone thioether. (C)1H-NMR spectra (offset ™0.5 ppm) showing the α-CH of the dimedone–thioether adduct at 4.70 ppm from cysteine and dimedone without (grey) and with SOD1 (blue) and compared to authentic CTS + dimedone (orange).

To detect sulfenic acids formed during the catalytic generation of thiosulfinates, cysteine oxidation by SOD1 was performed in the presence of dimedone (Fig. 2C) and compared with reactions lacking SOD1 (negative control) and with CTS-treated cysteine (positive control). ^1^H-NMR analysis revealed that SOD1-catalysed cysteine oxidation in the presence of dimedone produced a 1,3-cyclohexanedione thioether, mirroring the product profile observed for CTS driven cysteine sulfenylation (Fig. 2D). In contrast, cysteine and dimedone incubated without SOD1 showed no detectable conversion to the corresponding thioether. These data indicate that SOD1 generates thiosulfinates that undergo hydrolysis or nucleophilic attack by cysteine to sulfenic acids, which are trapped by dimedone.

### Reactivity of SOD1-generated Thiosulfinates

CBA oxidation by CTS is attributed to O-atom transfer (OAT) to the boronate reporter (29). The O-atom–donor capacity of CTS was assessed by comparing its reactivity with horseradish peroxidase (HRP) to that of H_2_O_2_. As expected, H_2_O_2_ quantitatively generated Compound I, producing the characteristic blue-shifted Soret band and increased absorbance at 500–650 nm (Fig. 4A) (30). Authentic CTS induced partial Compound I formation (Fig. 4A), indicating that CTS can generate high-valent heme intermediates, albeit less efficiently than H_2_O_2_. Despite this incomplete conversion, CTS supported robust HRP-dependent luminol oxidation (Fig. 4B), confirming that CTS behaves as an effective O-atom donor in an enzymatic context.

**Fig. 4.**
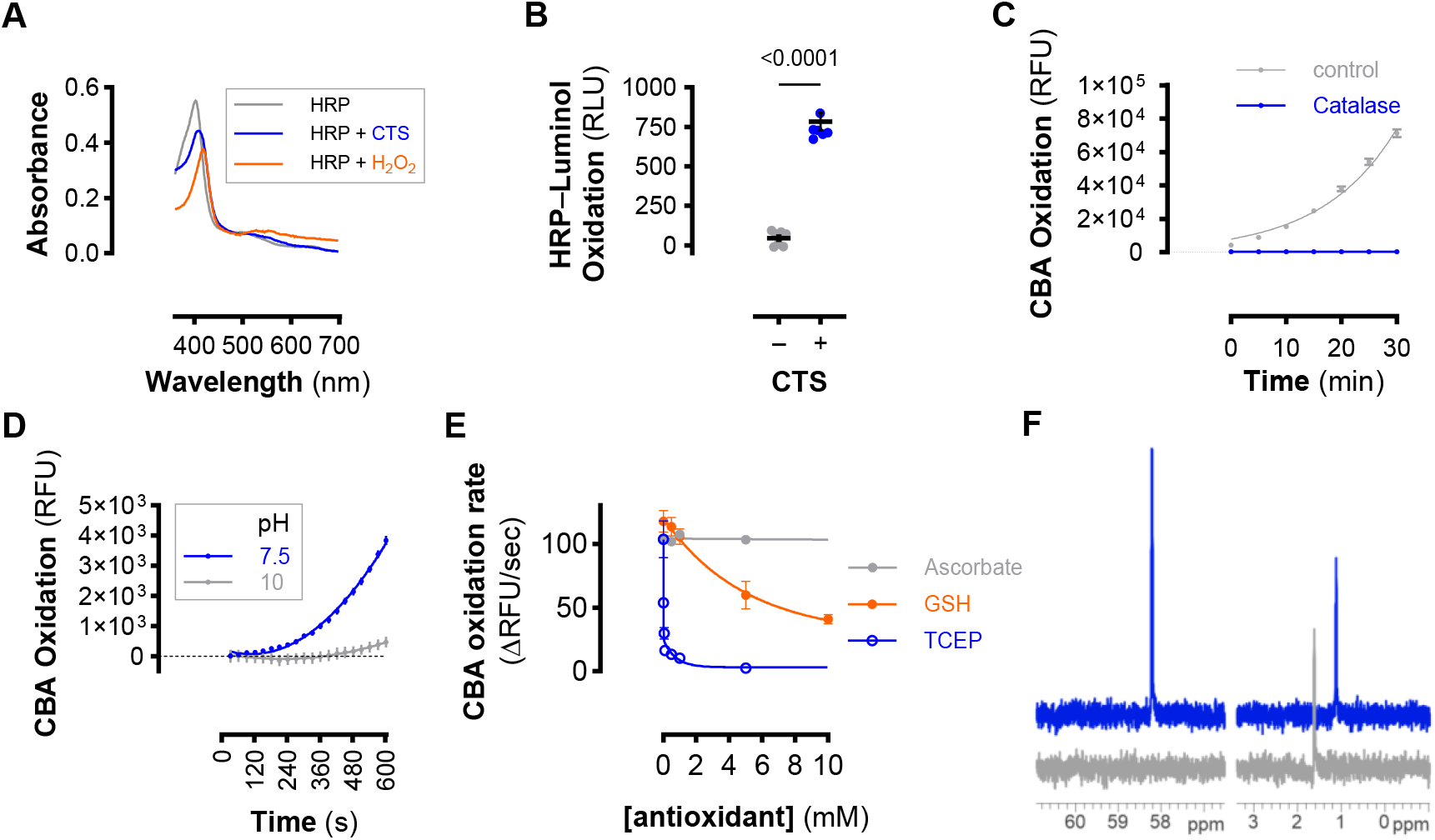
SOD1-generated CTS acts as a potent oxidant. (A). Visible spectra of HRP alone (grey) or reacted with CTS (blue) or H_2_O_2_ (orange). (B) Relative luminol luminescence (RLU) after reaction with HRP reacted with CTS (+) or vehicle (-). Lines represent mean ± SEM (n = 5) and analysed by unpaired, two-way t test. (C) The rate of CTS or CTS incubated with catalase induced CBA oxidation. Data are mean ± sd (n = 4) and nonlinear regression. (D) The rate of CBA oxidation by CTS at pH 7.5 (MOPS buffer) and 10 (borate buffer). Data points are mean ± SEM (n = 3) and fit to second-order curve (0.9969; 0.9836). (E) Effect of antioxidants on CBA oxidation. CBA (1 mM) oxidation rate from SOD1 (50 µM) and cysteine (300 µM) reaction in the presence of ascorbate, GSH or TCEP (0-10 mM). Data are mean ± SEM (n = 4) and GSH and TCEP data were fitted by nonlinear regression (two phase exponential decay model; r^2^_(GSH)_ = 0.794, r^2^_(TCEP)_ = 0.899). Ascorbate showed no concentration-dependent effect on CBA oxidation rates. (F) ^31^P-NMR spectra (offset ™0.5 ppm) showing formation of TCEP=O (+ 58.7 ppm) relative to phosphate (internal standard; +1.62 ppm) from TCEP in the presence of cysteine without (grey) or with SOD1 (blue).

CTS OAT reactivity was further validated using catalase pre-treatment, which abolished CTS-driven CBA oxidation (Fig. 4C), demonstrating that catalase consumes or intercepts the reactive O-atom equivalent. CTS-mediated CBA oxidation was also pH-sensitive, with activity nearly eliminated at pH 10 (Fig. 4D), consistent with loss of CTS-dependent OAT rather than cysteine sulfenic acid.

To probe antioxidant sensitivity of SOD1-catalysed CTS formation, the effects of ascorbate, GSH, and TCEP on CTS-dependent CBA oxidation were examined. Ascorbate produced no detectable inhibition up to 5 mM (Fig. 4E). To assess the relative kinetics of cellular thiol reactivity, GSH was titrated over a 0–10 mM range. Inhibition of CBA oxidation occurred only at high GSH concentrations, whereas equimolar GSH (1 mM) did not reduce CBA oxidation kinetics (Fig. 4E), indicating that GSH reacts with CTS much more slowly than CBA. TCEP potently inhibited CBA oxidation, yielding an IC_50_ of 9.6 µM, consistent with nucleophilic reduction of thiosulfinate. TCEP did not alter SOD1 activity (SI Appendix, Fig. S2E-F), indicating that inhibition reflects oxidant capture rather than enzyme inhibition. Accordingly, ^31^P-NMR detected formation of TCEP=O in SOD1–cysteine reactions (Fig. 4F). This confirms that SOD1-derived CTS functions as an O–atom donor.

### Physiological Relevance from Kinetic Modelling

Although SOD1 is canonically described as a superoxide-metabolizing enzyme, its active site is continuously exposed to abundant intracellular thiols. As free cytosolic O_2_^•-^ exists at sub- to low-nanomolar concentrations, and cysteine (Cys-SH) is present at ∼20–100 µM, SOD1 substrate availability differs by ∼10^5^–10^6^-fold. This disparity raises the possibility that thiol oxidation constitutes a physiologically relevant component of SOD1 reactivity, rather than an incidental reaction observed *in vitro*. Therefore, to estimate the potential contribution of thiol oxidation to overall SOD1 reactivity, pseudo–first-order rates were calculated using physiological substrate ranges and literature kinetics. Using the relationship v/[E] = (k_cat_/K_M_) [S], and literature values for SOD1-catalysed O_2_^•-^ (31) and cysteine kinetics derived here ((k_cat_/K_M_) O_2_^•-^ ≈ 2 × 10^9^ M^−1^ s^−1^ and (k_cat_/K_M_)Cys ≈ 3.32 × 10^2^ M^−1^ s^−1^, respectively), the rates of catalysis were compared across physiologic substrate ranges (ie, [O_2_^•-^] = 0.01–0.1 nM; [cysteine] = 10–100 µM; [homocysteine] = 0.1–1 µM) (32, 33). These calculations yield v/[E] ≈ 0.006–0.033 s^−1^ for cysteine oxidation and 0.02–0.2 s^−1^ for O_2_^•-^ dismutation within their respective basal ranges. Due to low physiological cellular levels, SOD1 metabolism of homocysteine is negligible. At resting cytosolic O_2_^•-^ levels, SOD1-catalysed cysteine oxidation therefore matches or exceeds the dismutation flux. A heatmap of these reactions shows that across the physiological cysteine range, thiol-oxidase turnover is competitive with canonical dismutation turnover (SI Appendix, Fig. S3A). A comparison of pseudo– first-order rates across representative substrate concentrations (SI Appendix, Fig. S3B) shows that SOD1-dependent cysteine oxidation occurs at rates overlapping those of O_2_^•-^ dismutation under physiological conditions. Within the physiological cysteine range, thiol-oxidase turnover can match basal dismutation flux, indicating that thiosulfinate formation is a major and continuous cellular activity of SOD1.

### Cellular Consequences of Thiosulfinate Formation

To define the cellular consequences of a bona fide thiosulfinate, HEK293 cells were exposed to increasing concentrations of cysteine thiosulfinate (CTS). CTS suppressed proliferation in a dose dependent manner and, at 500 µM, matched the antiproliferative effect of bolus H_2_O_2_ (Fig. 5A–B). CTS also depleted intracellular GSH and increased DCF fluorescence (Fig. 5C), consistent with strong oxidative activity. Because GSH does not react directly with SOD1 (Fig. 1B), but CTS is formed during SOD1 mediated cysteine oxidation and quenched by GSH (Fig. 4E), we examined whether GSH oxidation arises indirectly through CTS. Incubation of GSH with SOD1 in the presence of increasing cysteine produced a linear increase in GSH oxidation (Fig. 5D), consistent with cysteine dependent CTS formation driving downstream GSH consumption. Dimedone trapping indicated that CTS is the oxidant generated by SOD1; accordingly, the effect of dimedone on the SOD1 + cysteine reaction was tested. Dimedone (5–10 mM) inhibited CBA oxidation at pH 7.5 (Fig. 5E), consistent with interception of a thiosulfinate intermediate, and the requirement for excess dimedone reflects slower dimedone–CTS reaction kinetics relative to CTS driven CBA oxidation. Cellular oxidative stress mirrored these chemical reactivities. SOD1 alone did not induce DCF fluorescence, whereas SOD1 + cysteine and CTS did (Fig. 5F). Under the same conditions, dimedone based detection revealed robust protein sulfenylation, with SOD1 + cysteine and CTS producing similar Protein SOH profiles (Fig. 5G and SI Appendix, Fig. S4). Thus, CTS recapitulates the cellular oxidative signatures of SOD1 mediated thiol oxidation, supporting the conclusion that SOD1 generates a thiosulfinate like oxidant in cells.

**Fig. 5.**
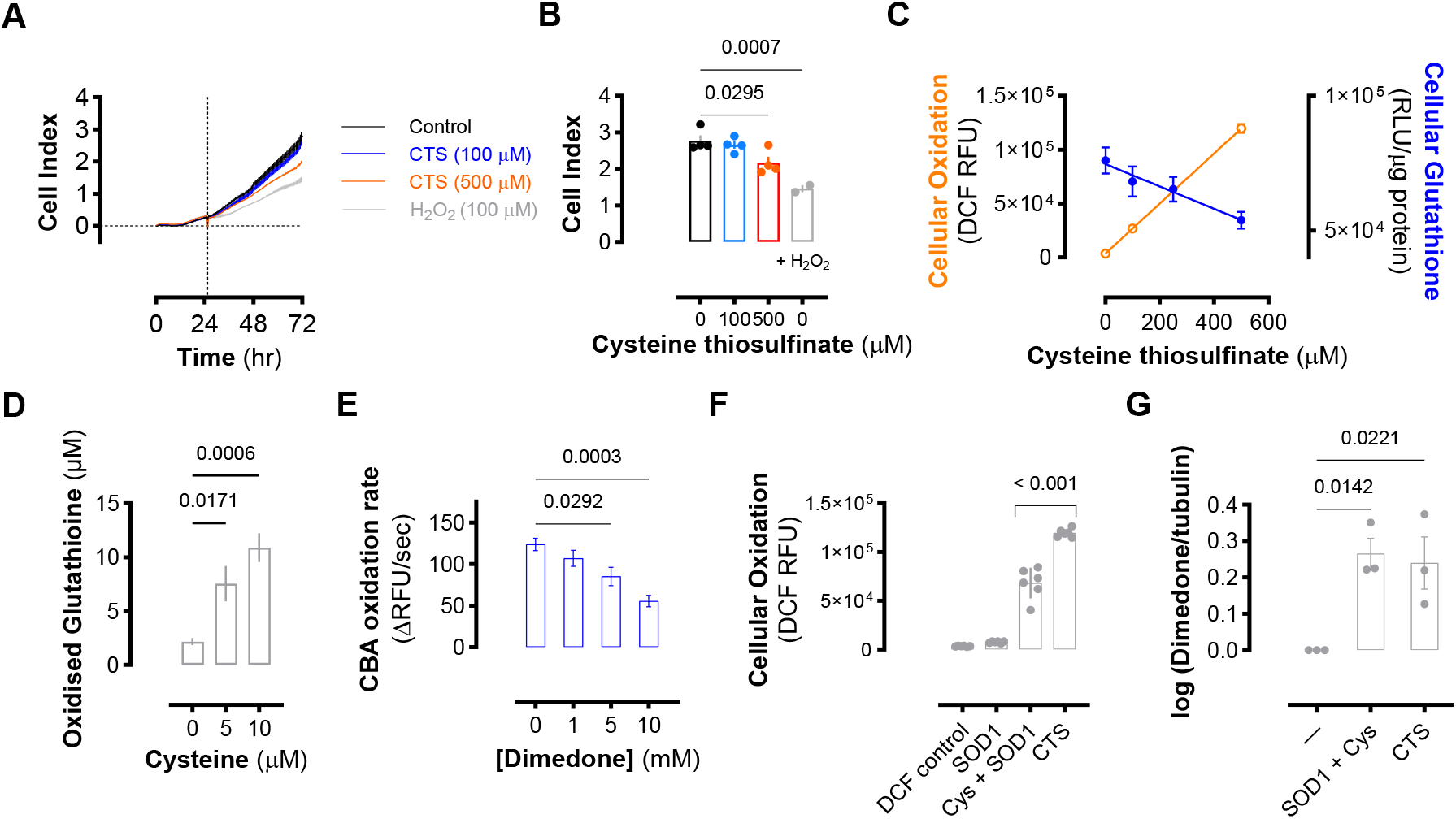
CTS phenocopies SOD1–thiol oxidation in cells. (A) Electrical-impedance measurement of HEK293 cellular proliferation treated with CTS or H_2_O_2_ and compared to untreated controls. CTS and H_2_O_2_ added to cells ≈ 24.5 hours represented by vertical dashed line. Data shown are mean ± sd (n = 2-4). (B) Bar graph showing the final cell index mean values ± sd (n = 2-4) from (A). Significance calculated by one-way ANOVA with Dunnett’s test. (C) Oxidative stress (orange, left axis) and intracellular glutathione (GSH) levels (blue, right axis) were quantified in HEK293 cells following treatment with authentic CTS. Oxidative stress was measured by DCF fluorescence and normalised to total protein content. Intracellular GSH was measured using a luminescent GSH-Glo assay, with relative light units (RLU) normalised to protein. Data represent mean ± SEM (n = 4-6 for both assays). Linear regression was applied to each dataset (r^2^ _(DCF)_ = 0.998; r^2^_(GSH)_ = 0.771). (D) Purified SOD1 was incubated with reduced glutathione (GSH) and increasing concentrations of cysteine to assess thiol oxidation. Following incubation, GSH oxidation was quantified using GSH-Glo luminescence. Data represent mean ± SEM (n = 5) and significance calculated by one-way ANOVA with Dunnett’s test. (D) CBA oxidation was measured in reactions containing SOD1 (50 µM) and cysteine (300 µM) in the presence of increasing concentrations of dimedone. Bars represent mean ± SEM (n = 5-7). Statistical significance was determined by one-way ANOVA with Dunnett’s test. (E) Bar graph showing relative oxidative stress via DCF fluorescence in HEK293 cells cultured with bovine SOD1 (50 µM), SOD1 + Cysteine (300 µM) or authentic CTS (500 µM). DCF RFU values were normalised to protein. Data represent mean ± SEM (n = 6) and significance calculated by one-way ANOVA relative to DCF controls. (F) Bar graph representing densitometric analysis of anti-dimedone-2-thioether immunoblots of HEK293 cells pre-treated with dimedone and then cultured with SOD1 and cysteine or authentic CTS. Dimedone labelling was normalised to tubulin and ratios were normalised to dimedone controls. Data shown as the log of dimedone/tubulin ratio and bars represent mean ± SEM (n = 3). Significance calculated by one-way ANOVA relative to dimedone controls.

To probe the cellular consequences of SOD1 thiosulfinate synthase activity, HEK293 cells were challenged with cysteine in pyruvate-free media (14). Under these conditions, cysteine (0.5–2 mM) caused a dose-dependent loss of viability. SOD1 inhibition with LCS-1 or ATN-224 significantly blunted this toxicity across the entire concentration range (Fig. 6A). SOD1 inhibition also suppressed homocysteine-induced cytotoxicity (Fig. 6B), indicating that SOD1 activity is required for thiol-dependent cytotoxicity.

**Fig. 6.**
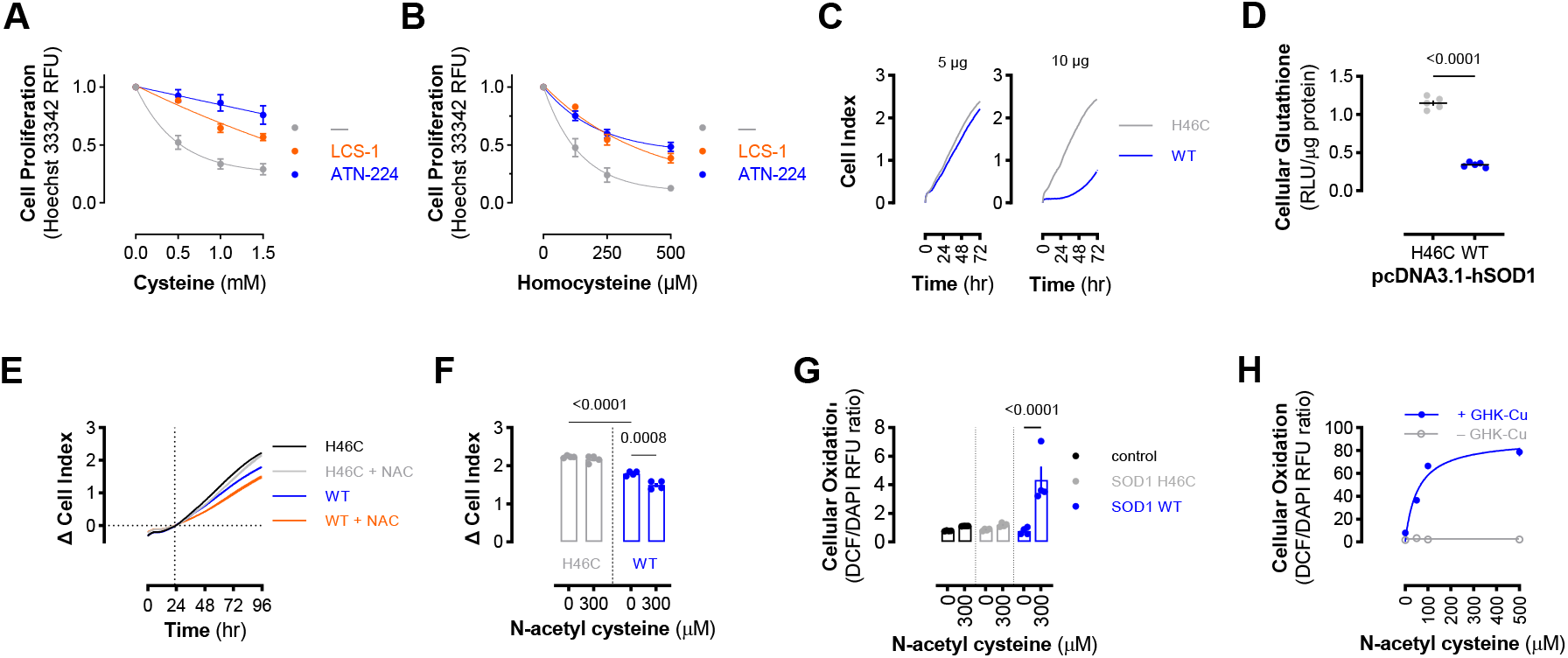
SOD1 activity mediates thiol-induced toxicity and oxidative stress. Proliferation of HEK293 cells in pyruvate-free MEM containing (A) cysteine or (B) homocysteine pre-treated with SOD1 inhibitors LCS-1 or ATN-224 and compared to vehicle (–) controls. Data are means SEM (n = 3) and fit to non-linear regression curves. (C) Representative electrical-impedance measurements of HEK293 cell proliferation overexpressing WT SOD1 (blue) or Cu-deficient (grey) SOD1 (H46C). (D) Relative cellular reduced glutathione (GSH) levels in HEK293 cells transfected (10 µg /1e^6^ cells) 48 hours post transfection. Relative luminescence units were normalised to protein. Lines represent mean ±SEM (n = 5) and significance calculated by unpaired, two-way t test. (E) Representative electrical-impedance measurement of proliferation for WT or H46C SOD1 overexpressing HEK293 cells treated with or without 300 µM N-acetyl cysteine (NAC). Differential cell index relative to time of NAC addition is shown. (F) Bar graph showing mean Δ cell index ±SEM (n = 4) of WT or H46C SOD1 overexpressing HEK293 cells ± NAC (300 µM). Significance calculated by two-way ANOVA. (G) Bar graph showing cellular oxidative stress in response to NAC treatment via DCF fluorescence normalised to DNA content in HEK293 cells transfected with WT or H46C SOD1 and compared to mock transfection controls. Bars represent mean ± SEM (n = 4) and significance calculated by two-way ANOVA. (H) Cellular oxidation as measured by DCF normalised to DNA content in HEK293 cells pre-treated with GHK-Cu or vehicle and NAC. Data are means ± sd (n = 3-6) and fitted to nonlinear regression (one phase exponential association model; r^2^_(+ GHK-Cu)_ = 0.918; r^2^_(-GHK-Cu)_ = −0.315).

To determine the contribution of SOD1-bound copper, WT SOD1 and the Cu-binding mutant H46C were compared. WT overexpression reduced proliferation and lowered intracellular GSH relative to H46C (Fig. 6C–D). NAC treatment further suppressed proliferation only in WT-expressing cells (Fig. 6E–F) and triggered a corresponding increase in DCF fluorescence (Fig. 6G), demonstrating that thiol addition selectively amplifies oxidative stress when active, copper-bound SOD1 is present. To test whether copper alone is sufficient to confer this phenotype, the SOD1-mimetic peptide GHK–Cu was evaluated. GHK–Cu reproduced the WT response, as NAC induced dose-dependent oxidative stress exclusively in GHK–Cu-treated cells and not in controls (Fig. 6H). Collectively, these results establish that copper-dependent SOD1 activity drives thiol-induced cytotoxicity, GSH depletion, and oxidative signaling through thiosulfinate formation, and that both cysteine-driven toxicity and NAC-sensitization arise from SOD1-derived thiol-oxidizing chemistry.

SOD1 expression is closely linked with cellular proliferation (6, 34, 35). siRNA-mediated SOD1 knockdown reduced growth relative to control cells (Fig. 7A–B and SI Appendix, Fig. S5) and was accompanied by a significant decrease in non-mitochondrial O_2_ consumption (Fig. 7C), consistent with SOD1 contributing to an O_2_-dependent reaction under physiological conditions. To test whether basal thiosulfinate formation accounts for this growth support, cells were treated with authentic CTS. Nanomolar levels of CTS enhanced proliferation in both control and SOD1-deficient cells and produced a hormesis-like response (Fig. 7D–E), while the cell-permeable thiosulfinate allicin yielded an even more pronounced hormetic window and similarly rescued SOD1 knockdown proliferation (Fig. 7F–G). These results show that basal SOD1-derived thiosulfinate formation supports cell growth at low thiol flux.

**Fig. 7.**
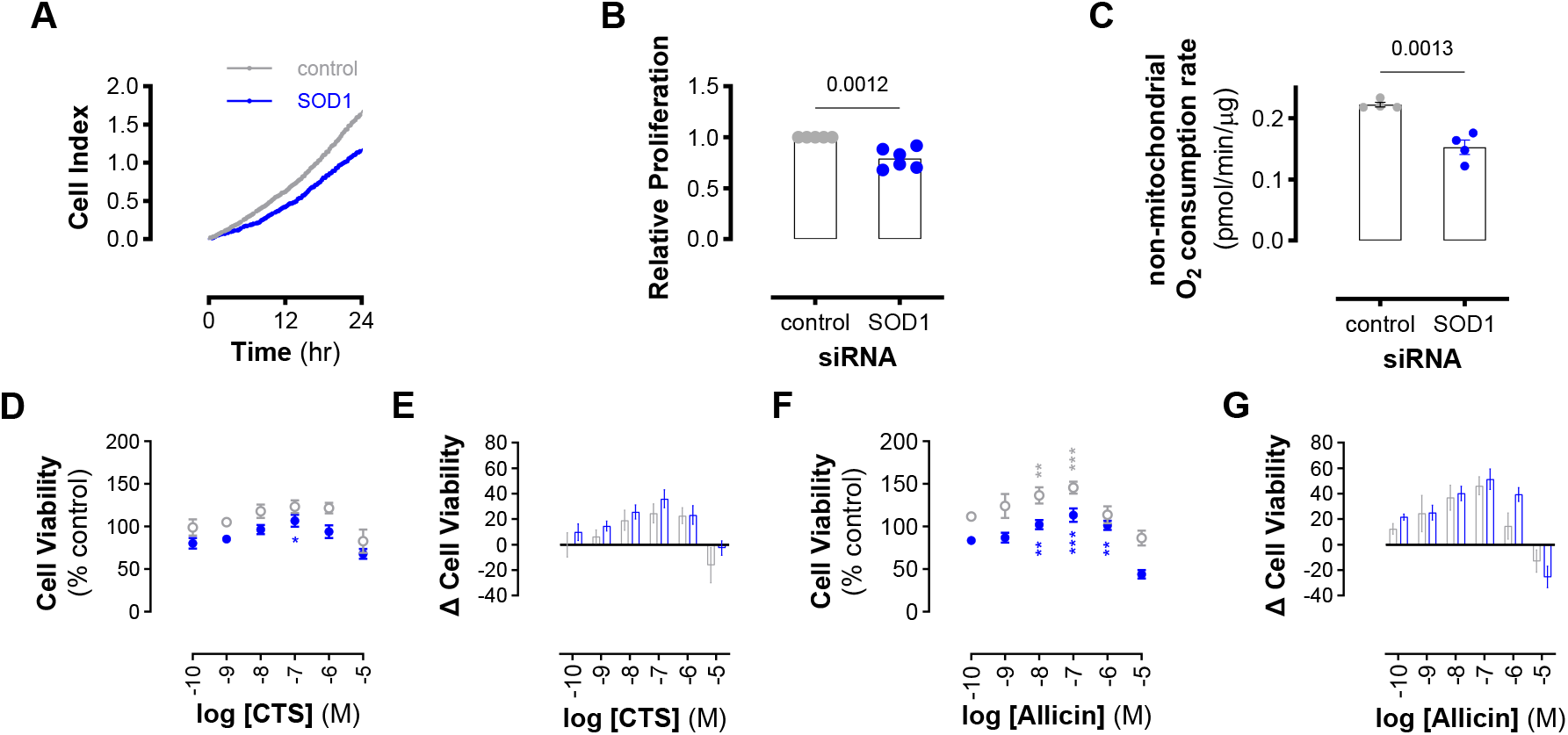
Basal SOD1 thiosulfinate activity supports proliferation. (A) Representative electrical-impedance measurement of HEK293 cell proliferation comparing control siRNA-treated cells (grey) and SOD1-knockdown cells (blue). Traces show real-time *Cell Index* values demonstrating reduced proliferation when SOD1 expression is decreased. (B) Relative proliferation of HEK293 cells after 24 h under control (grey) or SOD1-knockdown (blue) conditions. Bars show mean ±SEM (n = 6), normalised to control siRNA. (C) Non-mitochondrial O_2_ consumption in control (grey) and SOD1-knockdown (blue) cells. Data represent pmol O_2_ consumed per minute and normalised to protein content. Bars represent mean ± SEM (n = 4) and significance calculated from unpaired, two-way *t* test. (D) MTT-based viability measurements for control (grey open circles) or SOD1-knockdown (blue closed circle) HEK293 cells 24 hours after treatment with CTS. Data are mean ± SEM (n = 6) and normalised to % of untreated control cell viability. Significance calculated by one-way ANOVA with Dunnett’s test. (E) Bar chart showing the net differences in viability between CTS-treated and untreated cells for each phenotype (ie, Viability_CTS_ – Viability_control_). Data are mean ±SEM (n = 6). (F) Viability of HEK293 cells cultured with allicin. Data are mean ±SEM (n = 6) and normalised to % of untreated control cell viability. Significance calculated by one-way ANOVA with Dunnett’s test. (G) Net differences in viability between allicin-treated and untreated cells for each phenotype. Data are mean ±SEM (n = 6).

## Discussion

These results reveal an unanticipated dimension of SOD1 reactivity that broadens the landscape of cellular redox chemistry. Although SOD1 and low molecular weight thiols are traditionally framed as antioxidants, their interaction at the solvent exposed copper site (36) gives rise to a potent electrophilic oxidant capable of O-atom transfer (23, 37). Importantly, steady state analysis under physiological substrate ranges suggests that thiosulfinate formation can approach or rival the flux of superoxide metabolism, indicating that this activity is not a peripheral reaction but a significant component of SOD1 function within thiol rich cytosol. This places SOD1-mediated thiol oxidation alongside superoxide dismutation as a second major catalytic output of the enzyme.

The generation of a strong oxidant directly from thiols and molecular oxygen, without the need for canonical ROS intermediates, invites a broader conceptualization of redox biology in which sulfur based oxidative pathways operate in parallel with familiar ROS driven processes.

Formation of thiosulfinates may clarify longstanding inconsistencies in measurements of H_2_O_2_ during SOD1 reactions. Because thiosulfinates can drive oxygen atom transfer and sulfenylation that report as ‘peroxide’ in standard assays, their contribution would have inflated apparent superoxide dismutation signals, leading to systematic overestimation of superoxide derived H_2_O_2_ in earlier studies.

The continuous production of a reactive electrophile by an abundant cytosolic enzyme raises the possibility that SOD1 mediated thiosulfinate formation contributes to mitogenic signaling. Many thiol reactive oxidants and electrophiles exhibit hormetic behavior (38, 39), and the ubiquity of SOD1 together with the centrality of cysteine to metabolism suggests an evolutionary rationale for exploiting this chemistry during periods of oxygen availability. Under basal conditions, low flux thiosulfinate formation could provide a controlled oxidative signal that influences proliferation, consistent with the long recognized but mechanistically opaque growth promoting effects of SOD1 in diverse cellular contexts (40, 41).

When thiol concentrations rise beyond their physiological window, the same chemistry provides a unified explanation for the pronounced toxicity of both cysteine and homocysteine (16, 42).

Although their catalytic oxidation rates are similar, homocysteine may be more efficiently channeled into thiosulfinate formation because it has fewer competing metabolic fates, offering a mechanistic basis for its disproportionate cytotoxicity. This model resolves long standing uncertainties surrounding how elevated thiol levels trigger rapid oxidative stress despite the nominally antioxidant nature of the molecules involved (14, 22).

These principles have direct implications for tumor biology. Many cancers display concomitant upregulation of SOD1 (6, 43) and the cystine transporter SLC7A11 (44-46), creating a metabolic environment in which thiosulfinate formation could become a dominant oxidative pathway. Under such conditions, RS(O)SR species may act as potent mediators of oncogenic signaling or contribute to the heightened stress tolerance characteristic of cysteine addicted tumors (44-46). This perspective provides a chemical framework for understanding why interventions that elevate intracellular thiols, or that increase SOD1 abundance, can paradoxically enhance tumor aggressiveness.

Beyond cancer, altered thiosulfinate formation may be relevant in neurodegeneration, ageing, and other diseases in which SOD1 (wild type or mutant) is implicated (7, 11, 47, 48). The ability of SOD1 to generate electrophilic sulfur oxidants offers a plausible oxidative mechanism contributing to disease initiation or progression, independent of classical ROS pathways. This chemistry may represent a missing component in the ongoing debate over toxic gain of function mechanisms associated with SOD1, particularly in disorders where protein misfolding, redox imbalance, and disrupted thiol homeostasis converge (2, 7, 16, 18, 20).

A remaining challenge is the direct detection of thiosulfinates in living cells. Their intrinsic reactivity, rapid consumption by thiols, and conversion to non-diagnostic disulfides complicate direct observation, and their behavior often overlaps with that of peroxides in standard assays. Nonetheless, the demonstration that a highly abundant copper enzyme produces such oxidants from essential cellular thiols strongly suggests that thiosulfinates represent a biologically meaningful branch of sulfur metabolism.

In conclusion, these findings reposition SOD1 from a single purpose superoxide dismutase to a bifunctional oxidase that generates two mechanistically distinct oxidants: H_2_O_2_ and thiosulfinates. By revealing that SOD1 converts abundant cytosolic thiols into short lived sulfur electrophiles, this work broadens the chemical framework through which cellular redox signaling and stress responses are understood. Thiosulfinate formation provides a unified explanation for both the cytotoxicity of elevated thiol flux and the hormetic, pro-growth output of basal SOD1 activity, establishing sulfur-based oxidants as a second oxidative currency operating alongside ROS. Recognizing SOD1 as a modulator of both superoxide and thiol chemistry invites a reassessment of oxidative control across physiology, disease, and ageing, and highlights enzyme generated thiosulfinates as an overlooked but central component of cellular redox biology.

## Materials and Methods

Detailed experimental procedures are provided in SI Appendix. Briefly, SOD1-dependent thiol oxidation and oxidant formation were quantified using steady-state assays in defined buffer systems. Thiol consumption was measured by spectrophotometric and fluorometric methods, and oxidant formation was monitored using boronate-based probes. Time-resolved FT-IR spectroscopy and ^1^H-NMR were used to identify thiosulfinates and thiosulfinate-derived sulfenic acids. Oxidant reactivity assays included HRP Compound I formation, catalase-quenching controls, and nucleophile-competition experiments to assess O-atom transfer and electrophilic reactivity. Cell-based measurements were performed in HEK293 cells. Proliferation was assessed by real-time electrical impedance, intracellular oxidative stress by DCF fluorescence, and glutathione content by luminescent GSH assays. Protein-sulfenylation was quantified by dimedone-dependent immunoblotting. SOD1 activity was modulated through pharmacological inhibition, overexpression of WT or copper-deficient variants, and siRNA-mediated knockdown.

Kinetic modelling used experimentally determined rate constants and physiologic substrate ranges to compare thiol-oxidase turnover to superoxide dismutation under basal conditions. All data and additional methodological details are provided in the article and SI Appendix.

## Acknowledgments

The author acknowledges Prof. Phil Eaton for access to laboratory space during the early stages of this work and Ethan J. York for assistance with cell culture procedures. This work was supported by the Royal Society (RG\R1\241248).

## Author Contributions

C.H.S. designed the research, performed all experiments, analyzed the data, and wrote the paper.

## References

1. J. M. McCord, I. Fridovich, Superoxide dismutase. An enzymic function for erythrocuprein (hemocuprein). J Biol Chem 244, 6049–6055 (1969).

2. B. G. Trist, J. B. Hilton, D. J. Hare, P. J. Crouch, K. L. Double, Superoxide Dismutase 1 in Health and Disease: How a Frontline Antioxidant Becomes Neurotoxic. Angewandte Chemie International Edition 60, 9215–9246 (2021).

3. L. Papa, M. Hahn, E. L. Marsh, B. S. Evans, D. Germain, SOD2 to SOD1 Switch in Breast Cancer *. Journal of Biological Chemistry 289, 5412–5416 (2014).

4. S. Li et al., Disrupting SOD1 activity inhibits cell growth and enhances lipid accumulation in nasopharyngeal carcinoma. Cell Communication and Signaling 16, 28 (2018).

5. M. L. Gomez, N. Shah, T. C. Kenny, E. C. Jenkins, D. Germain, SOD1 is essential for oncogene-driven mammary tumor formation but dispensable for normal development and proliferation. Oncogene 38, 5751–5765 (2019).

6. S. Liu et al., SOD1 Promotes Cell Proliferation and Metastasis in Non-small Cell Lung Cancer via an miR-409-3p/SOD1/SETDB1 Epigenetic Regulatory Feedforward Loop. Frontiers in Cell and Developmental Biology Volume 8-2020 (2020).

7. L. Wang et al., Wild-type SOD1 overexpression accelerates disease onset of a G85R SOD1 mouse. Human Molecular Genetics 18, 1642–1651 (2009).

8. C. H. Switzer, S. Kasamatsu, H. Ihara, P. Eaton, SOD1 is an essential H(2)S detoxifying enzyme. Proc Natl Acad Sci U S A 120, e2205044120 (2023).

9. K. R. Olson et al., Metabolism of hydrogen sulfide (H2S) and Production of Reactive Sulfur Species (RSS) by superoxide dismutase. Redox Biology 15, 74–85 (2018).

10. C. C. Winterbourn, A. V. Peskin, H. N. Parsons-Mair, Thiol Oxidase Activity of Copper,Zinc Superoxide Dismutase*. Journal of Biological Chemistry 277, 1906–1911 (2002).

11. S. Bakavayev et al., Cu/Zn-superoxide dismutase and wild-type like fALS SOD1 mutants produce cytotoxic quantities of H(2)O(2) via cysteine-dependent redox short-circuit. Scientific reports 9, 10826–10826 (2019).

12. C. Montllor-Albalate et al., Sod1 integrates oxygen availability to redox regulate NADPH production and the thiol redoxome. Proceedings of the National Academy of Sciences 119, e2023328119 (2022).

13. M. D. Eaton, A. R. Scala, Toxic action of cysteine on ascites tumor cells in vitro. Experimental Cell Research 33, 481–494 (1964).

14. Y. Nishiuch, M. Sasaki, M. Nakayasu, A. Oikawa, Cytotoxicity of Cysteine in Culture Media. In Vitro 12, 635–638 (1976).

15. O. O. Pedersen, R. L. Karlsen, The toxic effect of L-cysteine on the rat retina. A morphological and biochemical study. Investigative Ophthalmology & Visual Science 19, 886–892 (1980).

16. C. E. Hughes et al., Cysteine Toxicity Drives Age-Related Mitochondrial Decline by Altering Iron Homeostasis. Cell 180, 296-310.e218 (2020).

17. M. Breau et al., The antioxidant N-acetylcysteine protects from lung emphysema but induces lung adenocarcinoma in mice. JCI Insight 4 (2019).

18. A. Hermann, G. Sitdikova, Homocysteine: Biochemistry, Molecular Biology and Role in Disease. 10.3390/biom11050737.

19. H. Jakubowski, Pathophysiological Consequences of Homocysteine Excess<sup>1</sup>. The Journal of Nutrition 136, 1741S–1749S (2006).

20. S. D’Elia et al., Homocysteine in the Cardiovascular Setting: What to Know, What to Do, and What Not to Do. 10.3390/jcdd12100383.

21. K. D. Held, J. E. Biaglow, Mechanisms for the oxygen radical-mediated toxicity of various thiol-containing compounds in cultured mammalian cells. Radiat Res 139, 15–23 (1994).

22. D. W. Jacobsen, Hyperhomocysteinemia and Oxidative Stress. Arteriosclerosis, Thrombosis, and Vascular Biology 20, 1182–1184 (2000).

23. E. Block, J. O’Connor, Chemistry of alkyl thiosulfinate esters. VII. Mechanistic studies and synthetic applications. Journal of the American Chemical Society 96, 3929–3944 (1974).

24. J.-H. Lee et al., Mechanisms of thiosulfinates from Allium tuberosum L.-induced apoptosis in HT-29 human colon cancer cells. Toxicology Letters 188, 142–147 (2009).

25. A. Roseblade, A. Ung, M. Bebawy, Synthesis and in vitro biological evaluation of thiosulfinate derivatives for the treatment of human multidrug-resistant breast cancer. Acta Pharmacologica Sinica 38, 1353–1368 (2017).

26. P. Nagy, M. T. Ashby, Reactive Sulfur Species: Kinetics and Mechanism of the Hydrolysis of Cysteine Thiosulfinate Ester. Chemical Research in Toxicology 20, 1364–1372 (2007).

27. Q. Zhu, C. Costentin, J. Stubbe, D. G. Nocera, Disulfide radical anion as a super-reductant in biology and photoredox chemistry. Chem Sci 14, 6876–6881 (2023).

28. N. M. R. McNeil, C. McDonnell, M. Hambrook, T. G. Back, Oxidation of Disulfides to Thiolsulfinates with Hydrogen Peroxide and a Cyclic Seleninate Ester Catalyst. 10.3390/molecules200610748.

29. J. Zielonka et al., Boronate Probes as Diagnostic Tools for Real Time Monitoring of Peroxynitrite and Hydroperoxides. Chemical Research in Toxicology 25, 1793–1799 (2012).

30. J. C. Toledo et al., Horseradish peroxidase compound I as a tool to investigate reactive protein-cysteine residues: from quantification to kinetics. Free Radical Biology and Medicine 50, 1032–1038 (2011).

31. J. A. Imlay, The molecular mechanisms and physiological consequences of oxidative stress: lessons from a model bacterium. Nature Reviews Microbiology 11, 443–454 (2013).

32. A. D’Angelo, J. Selhub, Homocysteine and Thrombotic Disease. Blood 90, 1–11 (1997).

33. M. P. Brigham, W. H. Stein, S. Moore, THE CONCENTRATIONS OF CYSTEINE AND CYSTINE IN HUMAN BLOOD PLASMA. The Journal of Clinical Investigation 39, 1633–1638 (1960).

34. C. K. Tsang et al., SOD1 Phosphorylation by mTORC1 Couples Nutrient Sensing and Redox Regulation. Molecular Cell 70, 502-515.e508 (2018).

35. X. Wang et al., SOD1 regulates ribosome biogenesis in KRAS mutant non-small cell lung cancer. Nature Communications 12, 2259 (2021).

36. J. A. Tainer, E. D. Getzoff, J. S. Richardson, D. C. Richardson, Structure and mechanism of copper, zinc superoxide dismutase. Nature 306, 284–287 (1983).

37. G. I. Giles, K. M. Tasker, C. Jacob, Oxidation of biological thiols by highly reactive disulfide-S-oxides. Gen Physiol Biophys 21, 65–72 (2002).

38. M. Nitti et al., Hormesis and Oxidative Distress: Pathophysiology of Reactive Oxygen Species and the Open Question of Antioxidant Modulation and Supplementation. 10.3390/antiox11081613.

39. S. Parvez, M. J. C. Long, J. R. Poganik, Y. Aye, Redox Signaling by Reactive Electrophiles and Oxidants. Chemical Reviews 118, 8798–8888 (2018).

40. X. Wang et al., Knockouts of SOD1 and GPX1 Exert Different Impacts on Murine Islet Function and Pancreatic Integrity. Antioxidants & Redox Signaling 14, 391–401 (2011).

41. K. Watanabe et al., Sod1 Loss Induces Intrinsic Superoxide Accumulation Leading to p53-Mediated Growth Arrest and Apoptosis. 10.3390/ijms140610998.

42. C.-G. Zou, R. Banerjee, Homocysteine and Redox Signaling. Antioxidants & Redox Signaling 7, 547–559 (2005).

43. S. König et al., Superoxide dismutase 1 mediates adaptation to the tumor microenvironment of glioma cells via mammalian target of rapamycin complex 1. Cell Death Discovery 10, 379 (2024).

44. Y. Okano et al., Oncogenic accumulation of cysteine promotes cancer cell-proliferation by regulating the translation of D-type cyclins. Journal of Biological Chemistry 300 (2024).

45. X. Tang et al., Cystine addiction of triple-negative breast cancer associated with EMT augmented death signaling. Oncogene 36, 4235–4242 (2017).

46. D. Tiek et al., Cysteine addiction in drug resistant glioblastoma and therapeutic targeting with designer selenium compounds. Neuro-Oncology 10.1093/neuonc/noaf265, noaf265 (2025).

47. G. Pharaoh et al., Metabolic and Stress Response Changes Precede Disease Onset in the Spinal Cord of Mutant SOD1 ALS Mice. Frontiers in Neuroscience 13 (2019).

48. A. Şahin et al., Human SOD1 ALS Mutations in a Drosophila Knock-In Model Cause Severe Phenotypes and Reveal Dosage-Sensitive Gain-and Loss-of-Function Components. Genetics 205, 707–723 (2017).

