## Supplemental Info for "SOD1 Catalyses Thiol Oxidation to Thiosulfinates"

Christopher H. Switzer

Christopher H. Switzer


**This PDF file includes:**

Supporting text
Figures S1 to S5
Table S1

**Supporting Information Text**

**Materials and Methods**

**1. Chemicals, reagents and assay kits**

Human and bovine erythrocyte SOD1, bovine liver catalase and horseradish peroxidase (HRP) were purchased from Sigma-Aldrich. L-cysteine, L-cystine, L-cysteine sulfinic acid, L-homocysteine, dihydrolipoic acid, glutathione, 3-chloroperbenzoic acid (≤77%), dimedone (5,5-dimethyl-1,3-cyclohexanedione), hydrogen peroxide ($H_2O_2$) (30%, ACS grade), LCS-1, luminol (sodium salt), potassium ferricyanide and 5,5-dithio-bis-(2-nitrobenzoic acid) (DTNB) were also obtained from Sigma-Aldrich. ATN-224, Coumarin Boronic Acid (CBA) and GHK-Cu were purchased from Cayman Chemical. BCA protein assay kit, 2′,7′-dichlorodihydrofluorescein diacetate (DCF-DA) and Hoechst 33342 (NucBlue) were obtained from Invitrogen. GSH-Glo™ Glutathione Assay was obtained from Promega. DMSO, MOPS, sodium acetate and sodium chloride were also from Sigma-Aldrich. Deuterated $d_6$-DMSO was obtained from Fisher Scientific. All buffers were prepared using ultrapure water and were further purified by filtration through 0.22 µm membranes before use.

Cysteine thiosulfinate (CTS) was synthesised according to literature procedure (23). Briefly, acidic L-cystine solutions were reacted with 3-chloroperbenzoic acid and CTS was precipitated, isolated by filtration, washed with methanol and dried overnight *in vacuo*. CTS was stored at -80C and used within 2 months. Purity of CTS was established by reaction with excess cysteine and calculating the equivalents of consumed cysteine via DTNB assay. CTS concentration was also verified before use by UV absorbance ($l_{max}$ = 247 nm) using an estimated extinction coefficient of ~ 150 $M^{-1}$ $cm^{-1}$ (47). Synthetic yields were routinely 80-90% of theoretical, with cystine the major contaminating species.

**2. SOD1 thiol oxidation**

**Thiol Consumption and Enzyme Kinetics**

Reactions were performed in 50 mM MOPS (pH 7.5) containing 150 mM NaCl at 25 °C, unless stated otherwise. Thiol substrates were mixed with human SOD1 at the indicated concentrations under pseudo–first-order conditions (thiol in excess over enzyme). At defined time intervals (typically every 5 min), aliquots were withdrawn and immediately quenched into excess DTNB. Absorbance at 412 nm was measured, and free thiol concentration was calculated. Thiol consumption was defined as the decrease in free thiol over time. Initial rates were obtained from the linear portion of the curve and plotted as a function of substrate concentration for Michaelis–Menten analysis. Nonlinear regression using GraphPad Prism v10 was used to determine $K_M$ and $k_{cat}$. Because the SOD1-catalysed reaction consumes two thiol equivalents per catalytic turnover, the apparent $k_{cat}$ was multiplied by 2 to reflect the overall stoichiometry.

**CBA Oxidation Assays**

Assays were performed in black, flat-bottom 96- or 384-well plates at 37°C in 50 mM MOPS (pH 7.5), 150 mM NaCl, with final concentrations human SOD1 = 10 µM and CBA = 1 mM. Reactions were initiated by automated injection of a L-cysteine stock solution (final 1 mM), and CBA fluorescence (coumarin product) was recorded kinetically at fixed excitation/emission settings using identical gain across conditions. Blank wells lacking enzyme and thiol were run in parallel and subtracted. To assess oxygen dependence, plates and cysteine solutions were equilibrated in the plate reader fitted with an atmospheric control unit set to 0.5% $O_2$ for 60 min prior to initiating reactions; kinetic acquisition proceeded under the same $O_2$ setpoint. To evaluate the effect of sulfenic acid trapping, reactions were performed as above with or without 10 mM dimedone; initial rates were compared under identical settings. Potassium ferricyanide was dissolved in MOPS buffer and added to reaction at 1 mM final concentration.

**pH Profiles for Thiol Consumption and CBA Oxidation**

pH-rate profiles were measured in 50 mM buffers containing 150 mM NaCl: sodium acetate (pH 6.5), MOPS (pH 7.5), and borate (pH 8.0 and 10.0). Standard conditions were SOD1 = 10 µM and L-cysteine = 1 mM; for CBA oxidation experiments, CBA = 1 mM was included. Reactions were initiated by cysteine addition, and initial rates were extracted as above to construct pH-activity profiles.

**ATR-FTIR Analysis**
The ATR detector was liquid-nitrogen-cooled and allowed to thermally stabilise before measurements. Authentic CTS. Authentic cysteine thiosulfinate (CTS) powder was applied directly to the ATR window. An air background was used as reference. Spectra were collected over 2000–900 $cm^{-1}$ using 16 co-added scans at 1 $cm^{-1}$ resolution. SOD1-cysteine reaction. Bovine SOD1 was prepared at 5 mg in 50 µL of 500 mM MOPS (pH 7.5), corresponding to a final enzyme concentration of 3.13 mM. A 300 mM L-cysteine solution was prepared fresh in the same buffer. To initiate the reaction, equal volumes (typically 2.5–5 µL) of bSOD1 and cysteine solutions were mixed directly on the ATR window. An immediate background spectrum was recorded, after which a repeated-scan kinetic acquisition protocol was initiated. Spectra were collected over the 1400–900 $cm^{-1}$ region at 1 $cm^{-1}$ resolution. Time-course spectra were acquired every 300 s for a total of six cycles. CTS-cysteine reaction. CTS was dissolved in 500 µL MOPS buffer to 0.3 mM final and immediately recorded by ATR-FTIR using buffer as the reference. To probe CTS reactivity with thiol, a 3 M cysteine stock was then added directly to the drop on the crystal to achieve 3 mM cysteine (final concentration) and spectra were acquired as for SOD1 reaction.

**NMR spectroscopy**
$^{1}$H-NMR spectroscopy
Dimedone trapping of sulfenic intermediates was assessed by $^{1}$H-NMR under three conditions: (i) L-cysteine (5 mM) + dimedone (25 mM) (enzyme-free control), (ii) bovine SOD1 (100 µM) + L-cysteine (5 mM) + dimedone (25 mM), and (iii) cysteine thiosulfinate (CTS, 5 mM) + dimedone (25 mM) (positive control). Reactions (total volume 500 µL) were carried out for 30 min at 37 °C in deuterated PBS (pD 7.8) and transferred immediately to 5 mm NMR tubes. Spectra were acquired at 298 K on a 400 MHz spectrometer using a standard 1D proton experiment (zg) with 32 scans, 2.5–3 s relaxation delay, and acquisition time 2–3 s. Chemical shifts were referenced to the residual HDO signal. The diagnostic α-CH proton of the dimedone–thioether adduct appeared at 4.70 ppm.

$^{31}$P–NMR spectroscopy
Phosphine oxidation was monitored by $^{31}$P-NMR. Cysteine (5 mM) and TCEP (5 mM) were prepared in 50 mM MOPS buffer (pH 7.5), and reactions were initiated by addition of either no enzyme, human SOD1 (20 µM), or SOD1 (20 µM) pre-incubated with ATN-224 (100 µM). Samples (500 µL) were incubated for 60 min at 37 °C, after which 100 µL $D_2O$ was added for field locking. Spectra were acquired at 298 K using a proton-decoupled $^{31}$P experiment (zgig), with 128–256 scans, a spectral width of 20–25 kHz, and a 2–3 s relaxation delay.

**SOD1 activity assay**
SOD1 activity was quantified using a xanthine/xanthine oxidase (X/XO)–driven WST reduction assay (Cayman Chemical Superoxide Dismutase Assay Kit), following the manufacturer's instructions. Briefly, reactions were assembled in the supplied assay buffer in 96-well format and contained WST working solution, xanthine substrate, and xanthine oxidase to generate superoxide at a constant rate. Human SOD1 (final concentration 2-200 nM) or TCEP (500 µM) was added to the reaction mixture immediately before initiating superoxide production with xanthine oxidase. WST reduction was monitored at 450 nm at 37 °C, and initial rates were calculated from the linear portion of the absorbance–time trace. Under these conditions, inclusion of TCEP did not alter SOD1-dependent inhibition of WST reduction.

**3. CTS Reactivity Assays**
CBA Reactivity with CTS and Catalase. Reactions were carried out in microplate format at 37°C. CTS was incubated with or without catalase for 10 minutes. Oxidation reactions were initiated by automated injection of CBA (1 mM final), and coumarin fluorescence was recorded every 5 min for 30 min (excitation = 332 nm; emission = 470 nm). Fluorescence traces were blank-subtracted.

CTS Reactivity with Cysteine. The reaction between CTS (3 mM) and L-cysteine (30 mM) was monitored spectrophotometrically by following the loss of CTS absorbance at 247 nm using a quartz microcuvette. CTS and cysteine were mixed at the stated concentrations in assay buffer, and $A_{247}$ was recorded at regular time intervals. CTS consumption was fitted using nonlinear pseudo-first order regression.

HRP Spectroscopy. HRP was first dissolved in water and diluted into 50 mM MOPS (pH 7.5) containing 150 mM NaCl. CTS was prepared in 50 mM acetate buffer (pH 5) and added as a 100× stock (1 µL into 100 µL final reaction volume). Final concentrations were HRP = 20 µM and CTS ≈ 70 µM. UV–vis spectra were recorded at room temperature using a CLARIOstar plate reader and compared with spectra obtained after addition of $H_2O_2$ (100 µM) or vehicle controls. HRP Luminol Reactivity. Horseradish peroxidase (HRP) was incubated with CTS or vehicle control under the same buffer conditions described above. Following the incubation period, luminol sodium salt was added to each reaction to a final concentration of 200 µM and allowed to react for 10 minutes at room temperature. Chemiluminescence was then quantified using a ClarioStar plate reader (BMG Labtech), and relative light units (RLU) were recorded.

### 4. Cell Culture, Transfection, and Proliferation

#### General Cell Culture

Human embryonic kidney 293 epithelial-like cell line (HEK293; ATCC) was maintained in Dulbecco’s Modified Eagle Medium (DMEM) (Gibco) supplemented with 10% fetal bovine serum (FBS) and Penicillin–Streptomycin–Neomycin (Invitrogen). Cells were cultured at 37 °C in a humidified incubator containing 5% $CO_2$ and were routinely passaged upon reaching 80–90% confluence.

#### SOD1 overexpression

HEK293 cells were transiently transfected with wild-type (WT) or H46C mutant SOD1 expression plasmids using Lipofectamine 3000 (Invitrogen) in Opti-MEM (Gibco), following the manufacturer’s instructions. The human SOD1 wild-type (WT) expression plasmid used in this study corresponds to the construct deposited at Addgene (plasmid #182922). The SOD1 H46C mutant expression plasmid was generated by GenScript using the CloneEZ ligation-independent cloning system and subsequently deposited at Addgene (plasmid # 253762). Both plasmids were sequence-verified prior to use. All plasmids were amplified in *E. coli*, purified using endotoxin-free maxiprep kits (Qiagen), and verified by Sanger sequencing prior to transfection. For experiments performed in 6-well plates (approximately $1 \times 10^6$ cells per well), 0, 5, or 10 µg of plasmid DNA was used per well. After 24 hours, cells were harvested for downstream assays, including xCELLigence impedance-based proliferation measurements and DCF-DA oxidation assays. Where indicated, N-acetylcysteine (NAC) treatments were applied by adding freshly prepared 100-fold stock solutions (*eg*, 1 µL stock into 100 µL medium) at the specified time points during the assay.

#### SOD1 knockdown

HEK293 cells cultured in T25 flasks or 6-well plates were transiently transfected with either Silencer Select negative-control No. 1 siRNA or Silencer Select SOD1-targeting siRNA (Thermo Fisher, assay ID s451) using Lipofectamine 3000 (Invitrogen) in Opti-MEM reduced-serum medium (Gibco), following the manufacturer’s protocol. After 24 hours, cells were either: seeded into 96-well plates for endpoint or fluorescence-based assays, seeded into E-plates for impedance-based proliferation measurements, or harvested and lysed for immunoblotting to confirm SOD1 knockdown.

#### Real-Time Cell Proliferation

Real-time cellular proliferation was monitored using the xCELLigence RTCA DP system (Agilent). E-plates were equilibrated with assay medium, after which cells were seeded at the desired density and placed in the RTCA station housed within a humidified 37 °C, 5% $CO_2$ incubator. Cell

Index (impedance-based proliferation readout) was recorded every 15 minutes throughout the experiment, beginning immediately after cell seeding and continuing before and after test compound addition.

**Thiol Toxicity Assays**

HEK293 cells were grown to approximately 80% confluence, washed briefly with pre-warmed medium, and overlaid with pyruvate-free Eagle's MEM (Gibco). Cells were pre-treated with the SOD1 inhibitors LCS-1 (10 µM) and ATN-224 (10 µM) for 1 hour prior to thiol exposure. Fresh solutions of L-cysteine or L-homocysteine were prepared in ultra-pure water immediately before use and added to cells as 100× stock solutions (1 µL stock per 100 µL final volume per well). After 24 hours, cell viability was assessed by quantifying DNA content using NucBlue™ Live ReadyProbes™ Hoechst 33342 staining. This DNA-based method was selected because elevated thiol concentrations produced high background signals that interfered with MTT-based metabolic and BCA protein assays. Fluorescence measurements were used to quantify viable cell number, and viability was normalized to cysteine-untreated controls within each inhibitor condition.

**DCF-DA Oxidative Stress Assays**

Intracellular oxidative stress was measured using 2',7'-dichlorodihydrofluorescein diacetate (DCF-DA), with appropriate precautions due to the dye's known susceptibility to artefacts, including light sensitivity, non-specific oxidation, and false-positive fluorescence signals. To minimize these issues, cells were incubated with DCF-DA under light-protected conditions, after which the dye-containing medium was removed and replaced with fresh assay medium. Cells were maintained in darkness throughout all subsequent steps.

Following dye loading, cells were treated with the indicated reagents to evaluate stimulus-induced ROS generation. Test reagents included bovine SOD1 (50 µM), cysteine (300 µM), cysteine sulfinic acid (300 µM), MMTS (300 µM), $H_2O_2$ (100 µM), and CTS (400 µM). Fluorescence was measured at 485/535 nm using a microplate reader at 37 °C. DCF fluorescence values were normalized to total protein content, quantified from assayed lysates using the bicinchoninic acid (BCA) assay. Background-subtracted values were normalized to untreated controls or as specified for individual experiments.

For CTS cell treatments, fresh CTS stock solutions were prepared in 50 mM acetate buffer (pH 5), diluted into culture medium, and applied to DCF-DA–loaded cells for 1 hour at 37 °C. Following fluorescence measurement, media were aspirated and cellular GSH was quantified as described in the corresponding section.

For HEK293 cells transfected with SOD1 WT or SOD1 H46C, Hoechst 33342 was co-loaded with DCF-DA to enable normalization to nuclear staining. A freshly prepared N-acetylcysteine (NAC) stock in PBS was added to culture medium to achieve a final concentration of 300 µM, and cells were incubated for 1 hour at 37 °C. DCF fluorescence values were normalized to Hoechst fluorescence to account for differences in cell number.
For GHK-Cu experiments, cells were similarly loaded with DCF-DA and Hoechst 33342, after which GHK-Cu (300 µM) and/or NAC (0–500 µM) were added to culture medium and incubated for 1 hour at 37 °C. DCF fluorescence was again normalized to Hoechst fluorescence.

**Cellular GSH assays**

Cellular reduced glutathione (GSH) levels were quantified using the GSH-Glo™ Glutathione Assay (Promega) following the manufacturer's instructions. Briefly, cells were lysed in the supplied lysis/reaction buffer, and lysates were incubated with the GSH-Glo reagent to enable conversion of the luciferin derivative proportional to the GSH content. Luminescence was measured using a microplate luminometer, and GSH concentrations were determined by interpolation from a freshly prepared GSH standard curve run in parallel. GSH values were normalized to total protein, quantified from matched lysates using the bicinchoninic acid (BCA) assay (Thermo Fisher Scientific).

**Non-mitochondrial oxygen consumption rates**

Non-mitochondrial oxygen consumption rates (OCR) were measured using an Agilent Seahorse XF extracellular flux analyser. Cells were seeded into Seahorse XF cell culture microplates at a density optimized for linear OCR response and allowed to equilibrate in Seahorse assay medium (supplemented as appropriate for the experiment) prior to measurement. Basal OCR was recorded, after which mitochondrial inhibitors (oligomycin, FCCP, antimycin A/rotenone) were sequentially injected following the manufacturer's recommended protocol to enable resolution of mitochondrial and non-mitochondrial components of $O_2$ consumption. Non-mitochondrial OCR was defined as the residual oxygen consumption rate following antimycin A/rotenone addition. At the end of each assay, plates were washed with ice-cold PBS, lysed, and total protein was quantified from matched wells using a BCA protein assay (Thermo Fisher Scientific). OCR values were normalized to micrograms of total protein and expressed as pmol $O_2$ $min^{-1}$ $\mu g^{-1}$ protein.

**Hormesis assays**

HEK293 cells were transfected with either control siRNA or SOD1-targeting siRNA using standard transfection conditions. Following transfection, cells were seeded into 96-well plates and allowed to grow for 12–24 hours until they reached approximately 70–80% confluence. Cells were then treated with thiosulfinate compounds, either CTS (dissolved freshly in acetate buffer, pH 5) or allicin (prepared as a concentrated stock in DMSO and subsequently diluted into cell culture medium to achieve final working concentrations). After compound addition, cells were incubated for an additional 24 hours under standard culture conditions. Cell proliferation was quantified using the CellTiter-Blue® Cell Viability Assay (Promega), following the manufacturer's protocol. Fluorescence was measured at 560/590 nm, and values were normalized to untreated control wells (no thiosulfinate).

**Immunoblotting**

Cells were lysed in RIPA buffer supplemented with protease inhibitors, and equal protein amounts were resolved on Bio-Rad 4–20% Tris SDS–PAGE gels. Proteins were transferred to PVDF membranes using a Bio-Rad Trans-Blot semidry transfer system. Membranes were blocked in 5% BSA in PBS–Tween (PBS-T) and incubated with primary antibodies overnight at 4 °C. The following primary antibodies were used: anti-SOD1 (1:1000; Cell Signaling Technology), anti-tubulin (1:2000; CST), and anti-dimedone (ABS30) (1:2000; Sigma-Aldrich). After washing in PBS-T, membranes were incubated with the appropriate HRP-conjugated anti-rabbit or anti-mouse IgG secondary antibodies for 1 hour at room temperature. Signals were developed using ECL™ Blotting Reagents (Cytiva, RPN2109), and membranes were imaged using the iBright imaging system (Thermo Fisher Scientific). Densitometry analysis was performed in ImageJ.

**5. Statistical Analyses**

All statistical analyses were performed using GraphPad Prism 10. Data are presented as mean ± SEM unless otherwise stated. Comparisons between two groups were assessed using unpaired two-tailed t-tests. For experiments involving more than two groups or multiple treatment conditions, one-way or two-way ANOVA was used as appropriate, followed by post-hoc multiple-comparison tests as specified in the legends. A $P$ value $< 0.05$ was considered statistically significant.

**Figures**

**A**

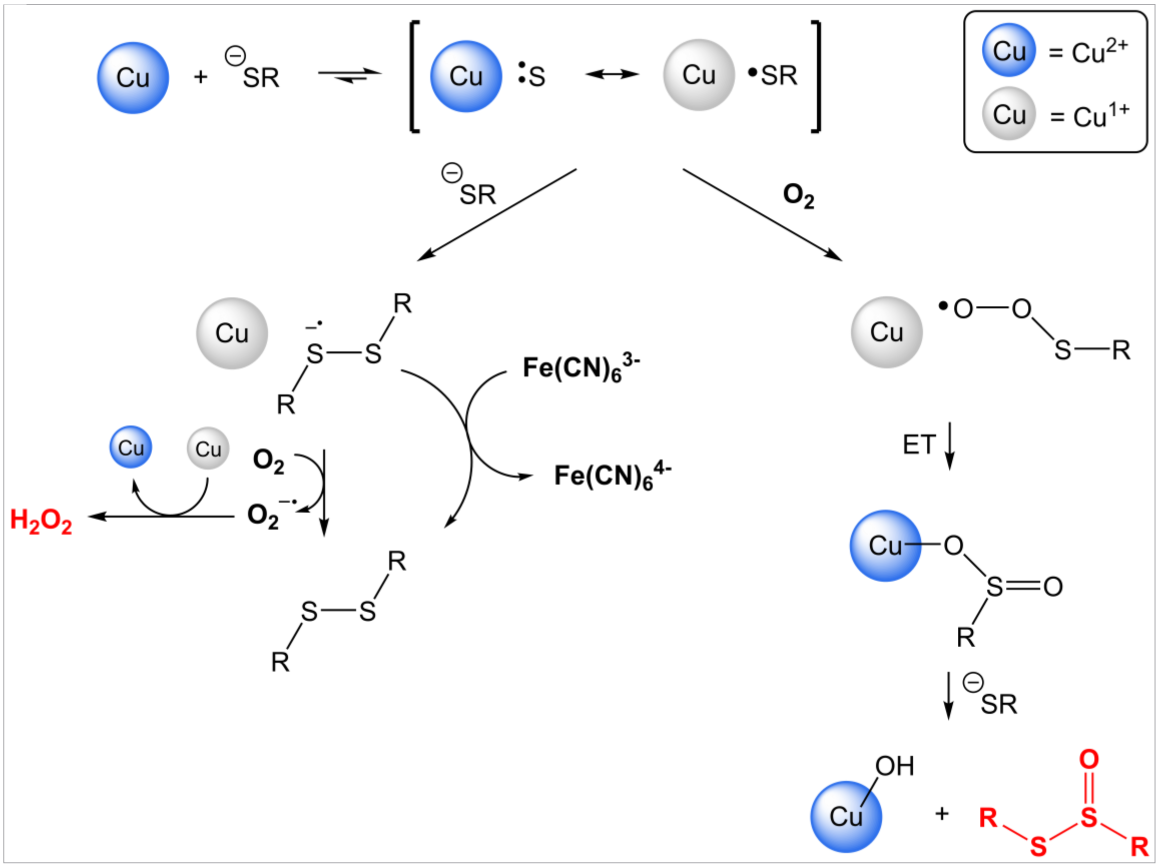


**B**

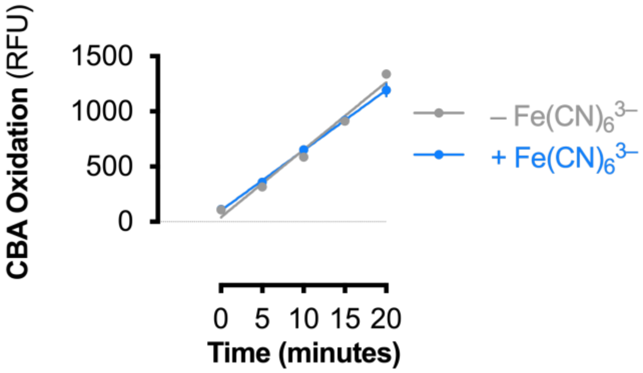


**Fig. S1. Mechanism of oxidant production from SOD1−thiol interaction.**
(A) Scheme of potential oxidant formation from SOD1 catalysed thiol reactions. Coordination of a thiolate to Cu(II)-SOD1 forms an intermediate with Cu(I)-thiyl character [Cu ·SR]$^+$. The radical intermediate can either react with another thiolate (RS-) to form a disulfide radical anion that reduces $O_2$ OR the [Cu ·SR]$^+$ radical intermediate can react with $O_2$ to form initially a Cu(II)−sulfinate and then, with attack of second thiolate, forms thiosulfinate [RS(O)SR].
(B) CBA oxidation over time by SOD1 and cysteine in the presence or absence of potassium ferricyanide ($K_3Fe(CN)_6$). Data are mean ± sd (n = 4) and fitted to linear regression analysis.

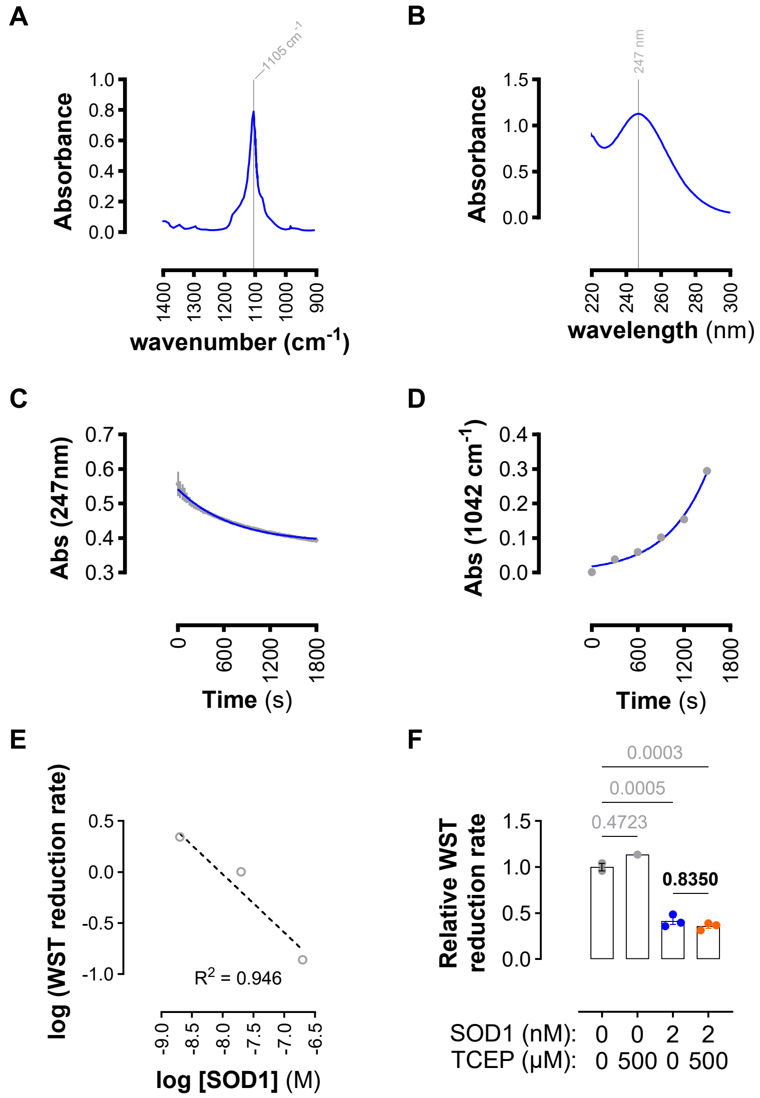


**Supporting Information Fig. S2 Authentic CTS characterisation and kinetics**

(A) ATR FT-IR spectrum of solid CTS with a local maximum at 1105 $cm^{-1}$.

(B) UV spectrum of CTS (approx. 1 mM) dissolved in 250 mM $H_2SO_4$ at 23°C with a local maximum at 247 nm.

(C) Rate of CTS consumption by cysteine. Data (grey) are mean ± SEM (n = 3) and fit to pseudo-first order curve (blue) (k = 0.001407 s-1; $r^2$ = 0.993).

(D) Rate of product formation from (A). Absorbance at 1042 cm-1 fit to pseudo-first order curve (blue) (k = 0.001881 s-1; $r^2$ = 0.990).

(E) Graph of SOD1 activity showing log (WST reduction rate) vs log [SOD1]. Data are mean ±sd (n =3) and linear regression analysis yielded $r^2$ = 0.948.

(F) Relative SOD1 superoxide dismutase activity in presence of TCEP. Data are mean ±sd (n = 2-3) and significance calculated by two-way ANOVA.

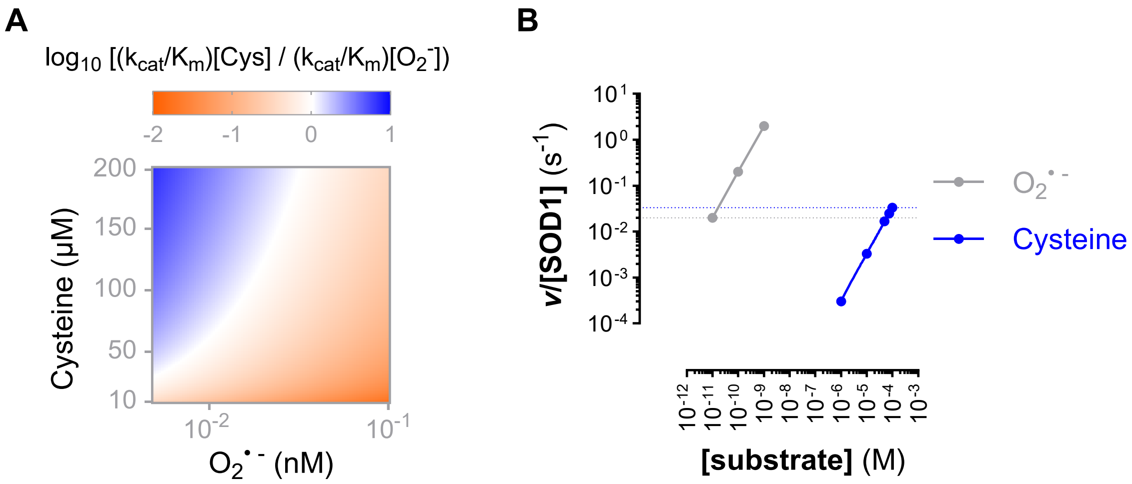


**Fig. S3. Comparison of SOD1 substrates: cysteine vs. superoxide ($O_2^{-}\cdot$).**
(A) Heatmap representing $\log_{10}[(k_{cat}/K_m)_{Cys}\cdot[Cys] / (k_{cat}/K_m)_{O2-\cdot}\cdot[O_2^{-}\cdot]]$. Blue regions indicate conditions where cysteine thiosulfinate rates exceed $O_2^{-}\cdot$ dismutation, whereas orange regions indicate $O_2^{-}\cdot$ dismutation rates dominate over thiol oxidation. White regions are conditions where cysteine and superoxide turnover rates are equal.
(B) Experimentally measured turnover rates (v/[SOD1]) plotted as a function of physiological cellular substrate concentration. The data show that cysteine supports turnover rates comparable to those observed with $O_2^{-}\cdot$, demonstrating that SOD1-catalysed cysteine oxidation, and therefore CTS formation, is a kinetically competent and cellularly relevant SOD1 activity.

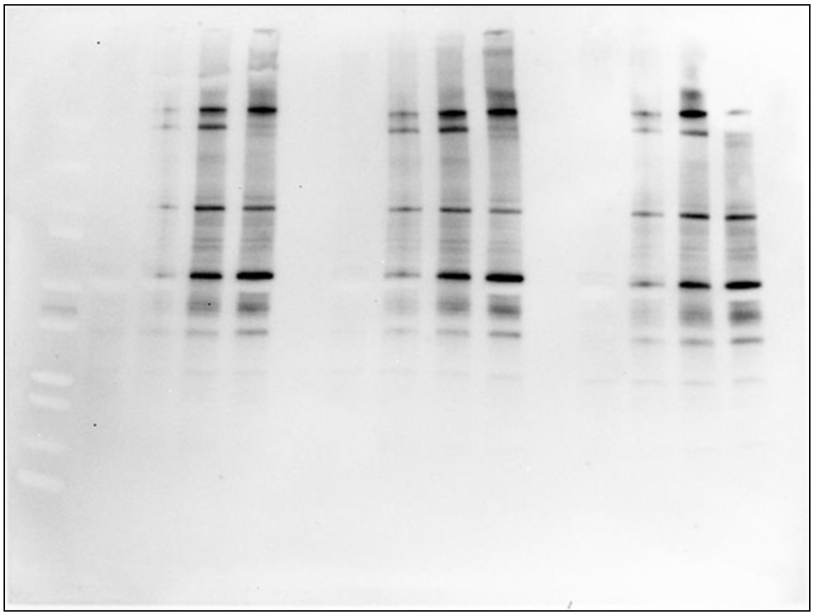

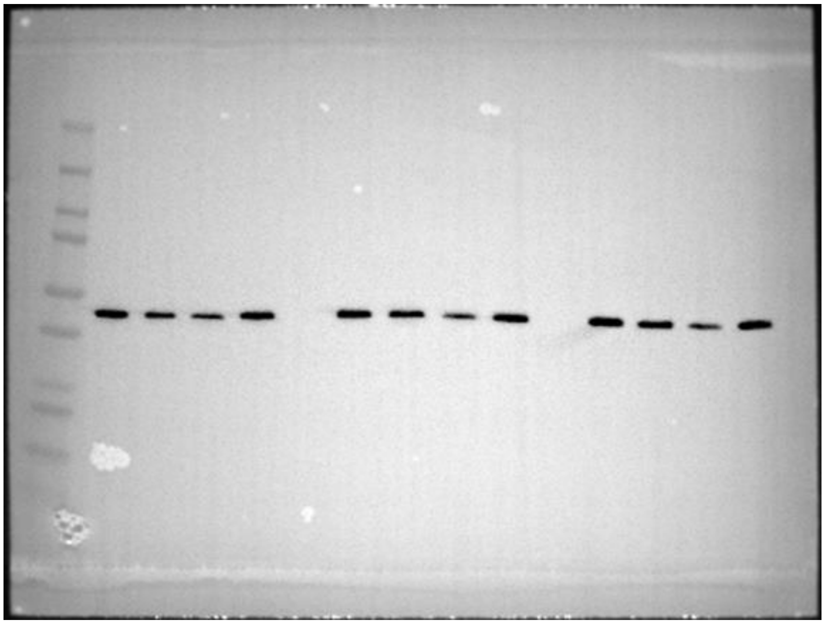

**Fig. S4. Protein-sulfenylation Immunoblot images**
Full immunoblots used for densitometric analyses in Fig. 5G for dimedone thioether (*top*) and (*bottom*) α-tubulin.

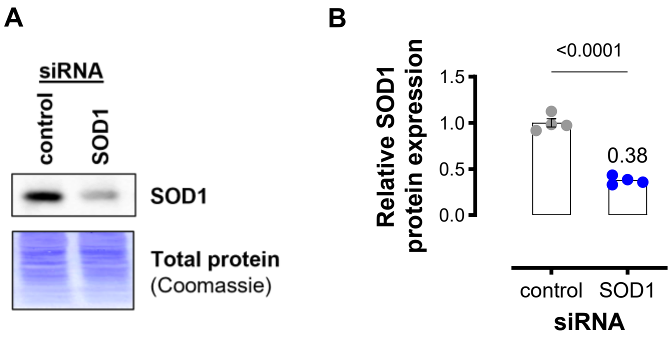


**Supporting Information Fig. S5. SOD1 knock-down quantification.**
(A) Immunoblot of SOD1 expression and total protein in representative HEK293 knock-down experiment used in Figs. 6-7.
(B) Bar graph showing relative SOD1 expression in control and SOD1 siRNA-treated HEK293 cells. Data are mean ± SEM (n = 4) SOD1 densitometric values normalised total protein (*au*) and are normalised to control mean.

**Tables**

**Table S1. SOD1-thiol Michaelis-Menten kinetics**

| | Vmax (s-1) | Km (mM) | R2 |
|---|---|---|---|
| L-cysteine | 5.94 | 3.57 | 0.999 |
| DL-homocysteine | 6.54 | 4.22 | 0.994 |
| $H_2$-LA | 2.00 | 4.63 | 0.994 |
| Glutathione | Unstable | 1.251E+15 | 0.959 |
